# Elevated conformational dynamics makes ACKR3 activation-prone and G protein-incompetent

**DOI:** 10.64898/2026.05.17.725760

**Authors:** Kai Wang, Tony Ngo, Ekta Khare, Rezvan Chitsazi, Suchismita Roy, Christopher T. Schafer, Tracy M. Handel, Irina Kufareva

## Abstract

The atypical receptor ACKR3 works together with the canonical chemokine receptor CXCR4 to drive cell migration along gradients of their shared agonist CXCL12. CXCR4 promotes chemotaxis by activating canonical G protein pathways and recruiting β-arrestins. ACKR3 indirectly regulates CXCR4-mediated chemotaxis by scavenging CXCL12. Unlike canonical chemokine receptors, ACKR3 does not couple to G proteins and instead is 100% biased towards β-arrestins. CXCR4 activation by CXCL12 is exquisitely sensitive to subtle changes in both receptor and ligand. By contrast, ACKR3 is activation-prone: it recruits β-arrestins in response to many ligands and is much less sensitive to mutations, suggesting distinct activation mechanisms compared to CXCR4. To explore the basis of these differences, we compared the dynamics of ACKR3 and CXCR4 complexes with chemokines using molecular dynamic (MD) simulations. Ten-microsecond atomistic MD simulations revealed that CXCR4 adopts a stable active state when bound to WT CXCL12 but transitions to an inactive state when in complex with the antagonist variant, [P2G]CXCL12. By comparison, ACKR3 exhibits a variable transmembrane (TM) 6 state distribution and persistently “active” TM7 when complexed with either WT CXCL12 or [P2G]CXCL12, the latter retaining substantial agonistic activity at ACKR3. We further identified ligand-mediated residue interaction networks in the TM core that regulate TM6 and TM7 activation in CXCR4 but are absent or disrupted in ACKR3, resulting in less constrained receptor dynamics. These findings were validated by BRET-based assays with CXCL12 and ACKR3 mutants. Together, the data suggests that the unique conformational dynamics of ACKR3 govern its activation propensity, its ligand promiscuity, and its atypical effector coupling.

## Introduction

The chemokine receptor CXCR4 promotes cell migration in response to the chemokine CXCL12 during numerous processes including the development of the vascular, immune and central nervous system, cardiac and renal organogenesis, as well as angiogenesis, homing of hematopoietic stem cells, and tissue regeneration and repair (1–4). CXCR4 is also involved in disease, most notably cancer, where it is upregulated in many solid tumors and cells in the tumor microenvironment as well as in hematologic cancer cells, and contributes to metastasis, angiogenesis and survival of the malignant cells (5–7). CXCR4 often works in concert with ACKR3, an atypical receptor that also responds to CXCL12. However, whereas CXCR4 directly promotes cell migration by CXCL12-mediated activation of G proteins, ACKR3 indirectly facilitates cell migration by scavenging CXCL12 to regulate its extracellular concentration and prevent overactivation and downregulation of CXCR4 (8–11). ACKR3 scavenges CXCL12 by continuously and constitutively internalizing into the cell, transporting the chemokine for degradation, and recycling back to the cell surface (12–14). CXCR4 does not constitutively internalize or recycle; instead, upon agonist stimulation, it is primarily targeted for degradation. ACKR3 functions without coupling to G proteins, instead robustly recruiting β-arrestins when stimulated by agonists (15,16). ACKR3 has also been reported to be basally associated with β-arrestins (17,18) but its constitutive trafficking can occur in the absence of β-arrestins (19). These functional distinctions suggest that ACKR3 and CXCR4 have different activation mechanisms.

Distinct activation mechanisms are also suggested by the contrasting sensitivity of the two receptors to different ligands and mutations. CXCR4 responds exclusively to CXCL12, and activation is extremely sensitive to mutations in both the ligand and the receptor (20,21). For example, a single-point P2G mutation in the N-terminus of CXCL12 ([P2G]CXCL12) renders it a pharmacological antagonist of CXCR4 (22). Additionally, mutations at the base of the orthosteric binding pocket of the receptor inhibit its ability to signal (23). This is consistent with experimental, computational and structural studies which suggest that signal transmission from CXCL12 to G protein engagement involves a precise network of residue interactions through the CXCR4 transmembrane (TM) helices (23–26), including the canonical class A GPCR microswitch residues (27,28). By contrast, ACKR3 is activated not only by CXCL12, but also by variants with N-terminal mutations including [P2G]CXCL12 (29,30), the chemokine CXCL11 (15), as well as other proteins (adrenomedullin (31), BAM22 (32)) and opioid peptides (16). With few exceptions, ligands that engage the orthosteric binding pocket of ACKR3 are activating, which is consistent with a more non-specific “distortion”-based activation mechanism (33,34). Specifically, we envision that steric interactions from ligand occupation of the ACKR3 binding pocket leads to rearrangements of its TM helices in a way that enables phosphorylation by G protein-coupled receptor kinases (GRKs) and recruitment of β-arrestins, but not the precise geometry required for G protein coupling.

Cryo-EM (34) and single-molecule FRET (smFRET) (33) studies have shed light on the unique structure and dynamics of ACKR3 compared to CXCR4. Structures revealed significant disorder in the intracellular loops (ICLs) of ACKR3 complexes (34), while smFRET showed that ACKR3 is dynamic and conformationally heterogeneous, readily exchanging between four roughly equally populated conformational states, both in the presence and absence of ligands (33). By contrast, apo-CXCR4 showed a major inactive state population and a less populated active state, as well as less probable state-to-state transitions, indicative of a more conformationally constrained receptor that requires greater ligand binding energy for activation (33). These insights into receptor dynamics are consistent with the propensity of ACKR3 to be activated by most ligands, and suggest that dynamic disorder in the intracellular pocket may preclude canonical effector coupling. However, the smFRET studies did not provide detailed mechanistic insights beyond global assessment of relative movements of the intracellular ends of receptor TM4 and TM6.

In this work, we explore high-resolution details of ACKR3 and CXCR4 activation via molecular dynamics (MD) simulations and pharmacological experiments. We first demonstrate that the atypical behavior of ACKR3 cannot be solely attributed to specific extracellular and intracellular sequence segments. Using comparative MD simulations, we reveal precise TM structural rearrangements that differ between ACKR3 and CXCR4, and residue interaction networks involved in the activation of CXCR4 that are absent or disrupted in ACKR3. Findings are validated in pharmacological experiments with rationally designed ACKR3 and CXCL12 mutants. Our work provides valuable insight for understanding how ACKR3 functions as an atypical receptor. It may also inform the design of ACKR3 inhibitors, which has been challenging due to its activation-prone nature.

## Results

### The inability of ACKR3 to couple to G proteins is not due to specific intracellular or extracellular motifs

Ligand activation of class A 7TM receptors and their subsequent coupling to G proteins generally involves conserved ‘microswitches’ in the receptor TM helices, including the DRY motif in TM3, CWxP motif in TM6, the NPxxY motif in TM7, and the PIF motif in TM3-5-6 (28). ACKR3 possesses all of these motifs, but it is still unable to activate G proteins (15,16). While the sequence flanking its DRY motif at the cytoplasmic end of TM3 is unusual (DRYLSIT), mutation to the canonical sequence (DRYLAIV) does not confer ACKR3 with the ability to couple to G proteins (35). We therefore sought other potential reasons for its G protein incompetence.

ACKR3 shares 31.4% sequence identity with CXCR4 in the TM domains (26.8% identity over the full sequence). Sequence alignment of ACKR3 with CXCR4 (**Fig. S1**) indicated that these receptors are particularly divergent in the N terminus and extracellular loops (ECLs) involved in ligand binding, as well as in the ICLs, helix 8 (H8) and the C terminus, involved in intracellular effector coupling (**Fig. 1A**). We therefore focused on these segments to design ACKR3 chimeras that might be able to activate G proteins, which was used as a surrogate metric for canonical GPCR behavior.

**Figure 1.**
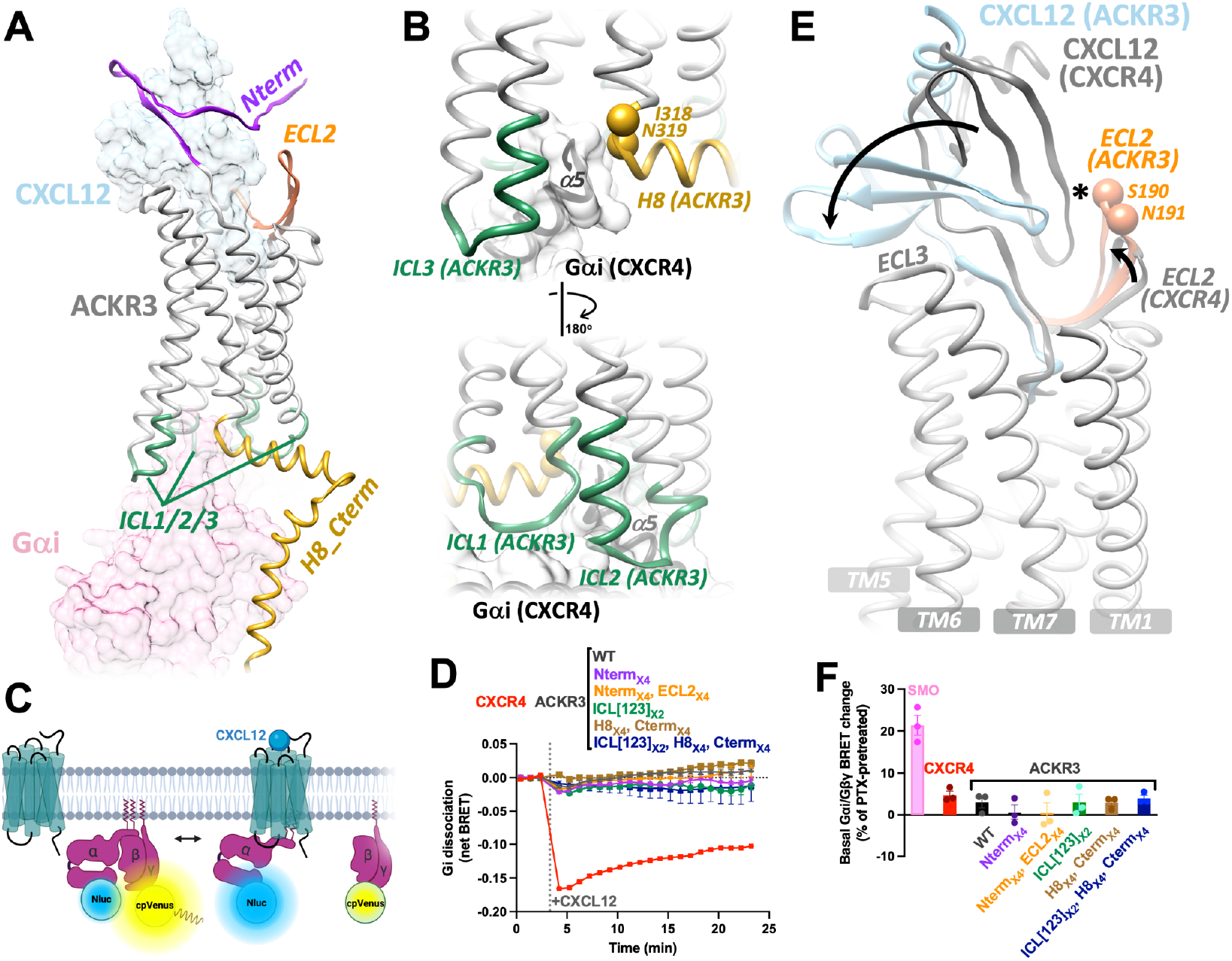
Chimeric ACKR3 variants containing extracellular and intracellular segments from CXCR2 and CXCR4 still fail to activate G proteins. (**A-B**) AlphaFold2 model of ACKR3:CXCL12 complex superimposed onto the cryo-EM structure of CXCR4 bound to Gi protein (PDB 8K3Z). Divergent extracellular (chemokine-interacting) and intracellular (effector-interacting) regions are highlighted in (A), while (B) shows the positioning of ACKR3 ICLs, the TM7-H8 hinge (Ile318-Asn319), and H8 at the Gαi “interface”. (**C**) Schematic of a BRET assay for the detection of CXCL12-stimulated Gαi and Gβγ subunit dissociation. (**D**) Representative net BRET time courses for CXCL12-induced Gαi/Gβγ protein dissociation assays with ACKR3 chimeras containing the canonical sequence substitutions for segments in (**A**). (**E**) ECL2 of ACKR3 is 2 residues longer than CXCR4, potentially preventing CXCL12 from forming a canonical orientation similar to that in CXCR4. “*”: clash between ACKR3 ECL2 and CXCR4-complexed CXCL12. (**F**) Basal Gαi/Gβγ protein dissociation measured by BRET relative to cells pretreated with pertussis toxin (PTX). Data are presented as mean ± SEM from three independent experiments.

Modeling based on our previously determined cryo-EM structures of ACKR3 suggested that clashes between ICL2 and/or ICL3 of the receptor with the N-terminal helix of Gαi could be responsible for the lack of G protein activity (34). However, chimeric ACKR3 variants containing individual or all ICLs from the canonical receptor, CXCR2, remained recalcitrant to G protein activation (34). Structural alignment of ACKR3 and CXCR4 also indicated that ACKR3 H8 (including the TM7-H8 hinge of residues Ile318-Asn319) could potentially interfere with the canonical core-insertion geometry of the α5 helix of Gαi (**Fig. 1B**). Therefore, building on prior findings from our lab (34) and the Stumm lab (35), we replaced the TM7-H8 hinge and H8 of ACKR3 together with its C terminus and/or all ICLs with the corresponding sequences from CXCR4 or CXCR2. The resulting chimeras still failed to activate Gαi proteins as assessed by a BRET-based G protein dissociation assay (**Fig. 1C-D**).

Cryo-EM structures (PDBs 7SK3 (34) and 8K3Z (36,37)) indicate that CXCL12 binds ACKR3 in a configuration that differs from CXCR4 as well as all other solved structures of receptor:chemokine complexes. This may be due to the divergent sequences of their N-termini and ECL2, which are directly involved in ligand engagement. For example, compared to CXCR4, the N-terminus of ACKR3 is 11 amino acids longer and has an altered distribution of acidic residues and reduced tyrosine content (**Fig. S2**), which may affect tyrosine sulfation patterns and chemokine engagement geometry (38,39). ECL2 of ACKR3 is two amino acids longer than CXCR4, which may also prevent CXCL12 from adopting an orientation similar to the CXCR4:CXCL12 complex (**Fig. 1E**). Therefore, we made chimeric variants of ACKR3 containing the CXCR4 N-terminus alone or together with ECL2, to determine if the changes conferred more canonical behavior. However, all these chimeras failed to activate Gαi proteins (**Fig. 1D, Fig. S3**).

All ACKR3 chimeric mutants were confirmed to properly express at the cell surface, ruling out aggregation or impaired trafficking as a cause of the lack of G protein activation (**Fig. S3**). We also confirmed that like WT ACKR3, all chimeras were devoid of basal Gi protein activity, by measuring basal Gαi/Gβγ dissociation with and without cell pretreatment with pertussis toxin (PTX) (**Fig. 1F**) (40). In contrast, PTX suppressed basal signaling of the constitutively active Smoothened receptor (SMO), used here as a positive control (40).

Together, these findings suggest that the non-conserved molecular composition of the extracellular and intracellular interfaces is not sufficient to account for the atypical pharmacology of ACKR3. Instead, we hypothesize that its atypical behavior is underlain by enhanced conformational dynamics, compared to canonical receptors like CXCR4, as suggested by cryo-EM structures (34) and smFRET studies (33). We thus turned to all-atom MD simulations to more broadly interrogate the conformational landscapes of these receptors and the structural determinants underlying their distinct dynamic behaviors.

### ACKR3 displays a dynamic TM6 and persistently “active” TM7, whereas CXCR4 exhibits ligand-dependent TM6/7 activation

To gain insight into the molecular mechanisms underlying the distinct conformational dynamics of ACKR3 and CXCR4, we generated all-atom MD simulations of the two receptors in complex with CXCL12 and [P2G]CXCL12, an N-terminal mutant of CXCL12 that is an agonist of ACKR3 but an antagonist of CXCR4 (21). AlphaFold2 (AF2 (41)) was used to model the extended receptor N-termini, which are disordered in cryo-EM density maps (34,36,37) but critical for chemokine binding and activation, in part by preventing chemokine dissociation (20,42). The AF2 models closely matched the subsequently solved experimental structures, with receptor RMSD_Cα_ below 0.8 Å and receptor:chemokine complex RMSD_Cα_ below 0.9 Å relative to the corresponding regions in the cryo-EM structures (PDBs 7SK3 and 8K3Z). Similar structural agreement was observed for predictions that were generated when AlphaFold3 (AF3 (43)) became available, supporting the reliability of predictions and utility of the computationally generated models for MD simulations (**Fig. S4**). Each of the above AF2 models was inserted in POPC bilayers, solvated, equilibrated, and simulated for 10 μs in triplicate to ensure unbiased sampling. Analyses of receptor tilt angles (**Fig. S5**), RMSD of TM domains (**Fig. S6**) and receptor:chemokine complexes (**Fig. S7**), as well as membrane properties (thickness (**Fig. S8**), area per lipid (**Fig. S9**), and lipid lateral diffusion (**Fig. S10**)) confirmed the stability of the complexes, appropriate membrane packing, and proper equilibration of the simulated systems. All simulations were initiated from active-state receptor conformations matching the cryo-EM structures. Consequently, the simulations probe the stability of the active state, and active-to-inactive state transitions, rather than activation events. Persistence of active-state features over the course of the simulation was interpreted as evidence of agonism, whereas their disruption was interpreted as receptor inactivation and antagonism, similar to observations in previous GPCR simulations (27,44).

To monitor global conformational changes of ACKR3 and CXCR4 in response to CXCL12 and [P2G]CXCL12, we plotted the distribution of the x- and y-coordinates of the intracellular ends of their TM helices over the course of three replicate simulations (**Fig. S11**). We initially focused on TM5, TM6, and TM7 (**Fig. 2**), the signature helices that undergo conformational rearrangements during GPCR activation (45,46). In two replicates (simulations 2 and 3), CXCL12-bound CXCR4 showed stable active-like positions of the intracellular ends of TM5/6/7 (indicated by limited x- and y-coordinate changes). In one replicate (simulation 1), CXCR4:CXCL12 transitioned to a more inactive state reflected by increased TM6 dynamics and an outward movement of TM7. This dynamic behavior is consistent with our prior smFRET study which showed that CXCL12-activated CXCR4 primarily populates a stable active state and to a lesser extent (∼31%), an inactive state conformation (33). In agreement with its antagonist pharmacology (21), the CXCR4 antagonist [P2G]CXCL12 triggered significant conformational fluctuations of TM6 (and TM5), and a more open conformation of TM7 in all three replicates, indicative of a transition towards an inactive state.

**Figure 2.**
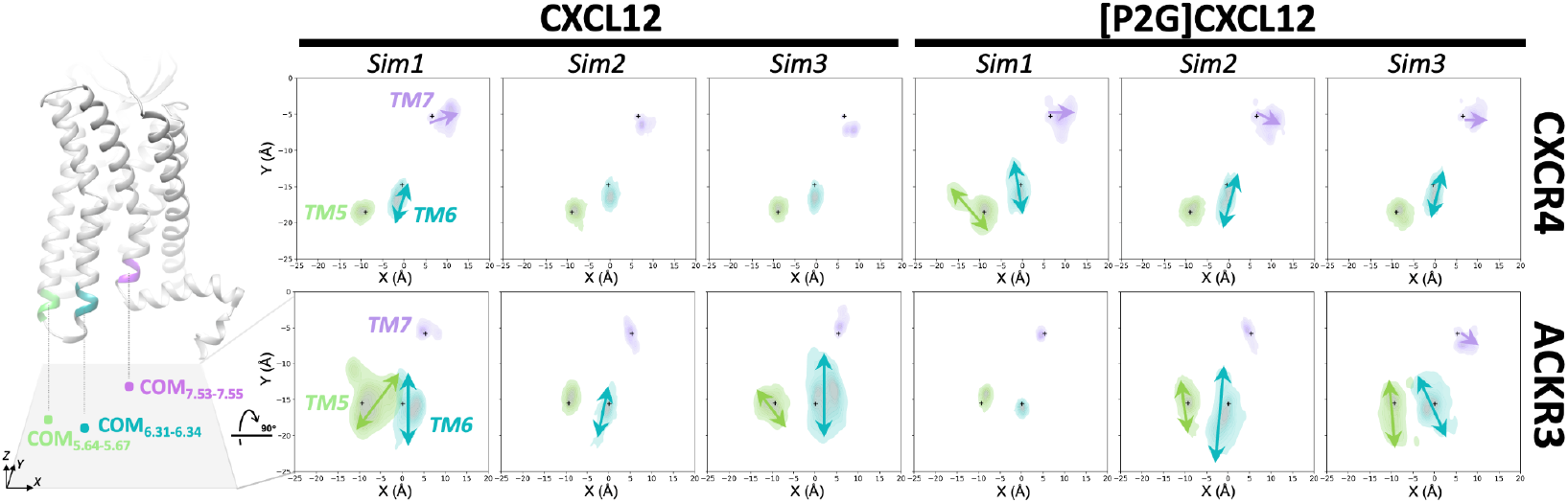
Dynamic TM5/6 and persistently “active” TM7 in ACKR3 contrast ligand-dependent TM rearrangements in CXCR4. Replicate 10 μs trajectories (n=3) aligned to receptor TM Cα atoms showing conformational dynamics of TM5, TM6, and TM7, represented as center-of-mass (COM) distributions on the xy-plane. Black “+” symbols indicate the starting conformations. Arrows indicate trajectories with large COM displacement relative to initial conformations; arrows corresponding to small displacements are omitted for clarity.

By contrast with CXCR4, ACKR3 showed a substantially more dynamic TM5 and TM6 in 5/6 simulations (all but simulation 1 of ACKR3:[P2G]CXCL12), regardless of whether it was occupied by CXCL12 or [P2G]CXCL12. This increased dynamics prevented ACKR3 TM5 and TM6 from maintaining a canonical active position. By contrast, a stable active-like conformation (inward positioning) of TM7 was observed in both CXCL12- and [P2G]CXCL12-complexed ACKR3 in 5/6 simulations (all except simulation 3 of ACKR3:[P2G]CXCL12). The indiscriminate conformational dynamics in response to CXCL12 vs [P2G]CXCL12 reflects a fundamental difference in ACKR3 compared to CXCR4, in chemokine signal initiation and/or transmission, and is consistent with ACKR3 activation by distortion rather than a specific network of interactions. As previously proposed, the increased dynamics or altered positioning of the ACKR3 TM helices could also lead to an inappropriately organized intracellular pocket that prevents G protein coupling (33,34).

### CXCL12 Leu5 is essential for canonical activation of CXCR4 but dispensable for ACKR3

Most known receptor:chemokine complexes share a global architecture where the chemokine globular domain interacts with the proximal receptor N-terminus (chemokine recognition site 1, CRS1) and ECLs, whereas the flexible chemokine N-terminus inserts into the receptor orthosteric pocket to engage TM residues (CRS2) and trigger receptor activation (47–49). CXCR4 and ACKR3 complexes with CXCL12 also abide by this architecture, but differ in chemokine orientation and CRS interactions. In CXCR4, CRS1 forms contacts with the CXCL12 N-loop, similar to other receptor:chemokine complexes, whereas in ACKR3, CRS1 interacts with the β1-strand of CXCL12 (**Fig. S12**). As a consequence, the CXCL12 globular domain is rotated by ∼80° in the ACKR3 complex relative to the CXCR4 complex; it is also shifted toward TM5 and TM6 in ACKR3, allowing it to interact primarily with ECL3 and the N-terminal end of TM5 rather than with the extended ECL2 observed in the CXCR4 complex (**Fig. 1E, Fig. S12**).

Substantial differences between the two receptors are also observed in CRS2. The N-terminus of CXCL12 inserts into orthosteric pockets of both CXCR4 and ACKR3 but is rotated ∼45° in ACKR3, compared with CXCR4, and forms a distinct network of interactions with the pocket (**Fig. 3A**). In CXCR4, consistent with prior modeling and structural studies (20,36,37,50,51), the N-terminal amine and the side chain of CXCL12 Lys1 form well-defined polar interactions with Asp97^2.63^, Asp171^4.60^, and Asp187^ECL2^ (superscripts indicate the universal Ballesteros-Weinstein (BW) numbering system for GPCRs (52)) (**Fig. S13**), both previously shown to be critical for CXCR4 activation (20,23,51,53–55). CXCL12 Pro2 packs against Trp94^2.60^, Tyr116^3.32^ and nearby pocket residues (**Fig. S13**), similarly important for CXCR4 signaling (20,23,36,37,50,51,53–56). The observed interactions explain why CXCL12 variants with deleted or mutated Lys1 and Pro2 are unable to activate CXCR4 (21,29,57). In ACKR3, CXCL12 Pro2 also packs against Phe124^3.32^ whereas the side chain of Lys1 is reoriented and interacts with Asp179^4.60^, Glu213^5.39^, Ser198^ECL2^, and Tyr200^ECL2^ (**Fig. S13**). However, neither mutations of these pocket residues (34,58–60) nor variations in the first four amino acids of CXCL12 (21,29,59) abrogate ligand-induced β-arrestin recruitment to ACKR3, consistent with a distinct activation mechanism.

**Figure 3.**
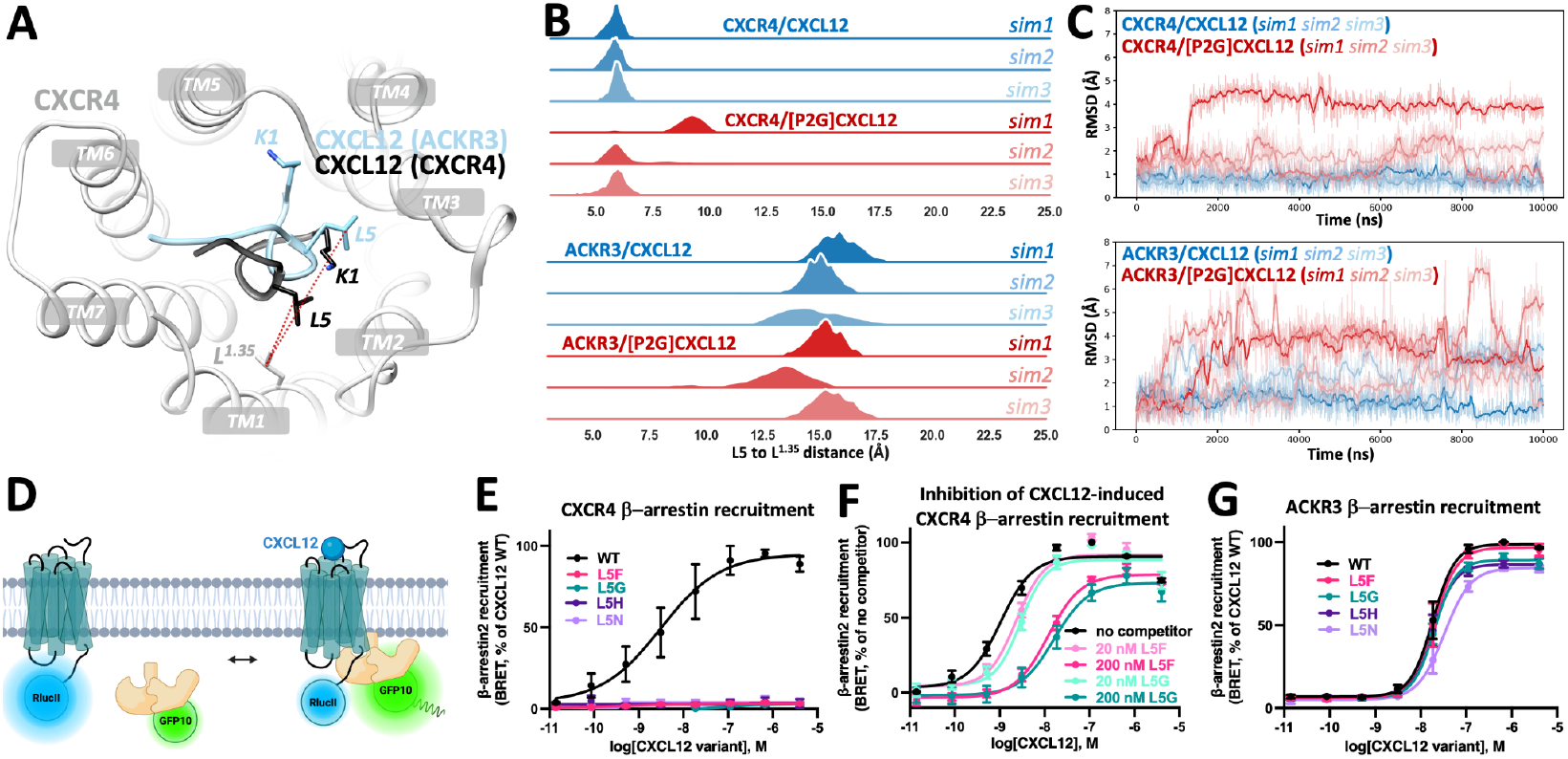
Structural comparison and functional analysis of CXCL12 Leu5 interactions with CXCR4 and ACKR3. (**A**) Structures of the chemokine-binding pockets in ACKR3 (PDB 7SK3) and CXCR4 (PDB 8K3Z) highlighting the orientation of the CXCL12 N-terminus in each receptor complex. (**B**) Distance distribution between the Cγ atoms of CXCL12 Leu5 and receptor residue Leu^1.35^, measured over replicate MD simulation trajectories of CXCR4 and ACKR3 complexes. The measured distances are indicated by dashed lines in panel (**A**). (**C**) RMSD of CXCL12 Leu5 relative to starting structures in complexes of CXCR4 or ACKR3 bound to WT CXCL12 or the mutant [P2G]CXCL12 over simulation time. Trajectories were aligned with receptor TM Cα atoms and RMSD was calculated with Leu Cγ atom positions. (**D**) Schematic of a BRET assay for the detection of CXCL12-stimulated β-arrestin2 association. (**E-G**) BRET in HEK293T cells co-expressing receptor-RlucII and GFP10-β-arrestin2 and stimulated with various concentrations of WT CXCL12 or CXCL12 variants (L5F, L5G, L5H, and L5N). Shown are CXCR4 activation (**E**), competition assay assessing CXCR4 engagement by CXCL12 mutants (**F**), and ACKR3 activation (**G**). Responses were expressed as % maximal response relative to WT CXCL12 with no competitor within each experiment and are presented as mean ± SEM from three independent experiments.

To better understand the divergent selectivity of the two receptors and how the CXCL12 N-terminus interacts with the orthosteric pockets of ACKR3 and CXCR4 to induce activation-related conformational changes, we examined the dynamics of the chemokine-CRS2 interactions over the course of the simulations. In addition to interactions involving Lys1 and Pro2, we noticed that Leu5 of CXCL12 forms a stable hydrophobic contact with Leu41^1.35^ (∼6 Å between the Cγ atoms) in CXCR4 (**Fig. 3B**). Consistent with the importance of this interaction, introduction of the P2G mutation in CXCL12 increased the dynamics of Leu5 in the CXCR4 complex (**Fig. 3C**), which destabilized its contact with Leu41^1.35^ (**Fig. 3B**). By contrast, in the ACKR3:CXCL12 complex, Lys1-Leu5 of CXCL12 are oriented toward TM3/4 and away from TM1/7, preventing the Leu5-Leu47^1.35^ interaction (>13 Å) (**Fig. 3A-B**). As a result, Leu5 exhibits higher baseline mobility, which is further increased by the P2G mutation (**Fig. 3C**). These observations suggest that the dynamic behavior of Leu5 contributes to the antagonistic activity of [P2G]CXCL12 toward CXCR4, and that the stable Leu5-Leu41^1.35^ interaction is important for canonical CXCR4 activation. In ACKR3, however, Leu5 appears less important due to the distinct chemokine-receptor geometry, consistent with the mutability of CXCL12 residues 1-4 (21,59).

To experimentally test this hypothesis, we mutated CXCL12 Leu5 to various amino acids with a broad range of sizes and chemical properties (Phe, Gly, His, and Asn) (61), and measured their ability to stimulate β-arrestin recruitment to CXCR4 and ACKR3 by BRET (**Fig. 3D**). All CXCL12 mutants abolished β-arrestin recruitment to CXCR4 (**Fig. 3E, Table S1**), confirming the critical role and precise chemical requirements for Leu5 in CXCR4 activation. The loss of agonism did not occur though the loss of binding, because the mutants were able to inhibit CXCR4 activation by WT CXCL12 in a concentration-dependent manner (**Fig. 3F**). By contrast with CXCR4, all Leu5 mutants retained agonism towards β-arrestin recruitment to ACKR3 (**Fig. 3G, Table S1**). Together, these results demonstrate that CXCL12 Leu5 is critical for canonical CXCR4 activation, but not for atypical ACKR3 activation, due in part to differences in binding modes and receptor conformational dynamics.

### ACKR3 lacks the intramolecular hydrophobic spine linking CXCL12 Leu5 to TM7 activation

To investigate how CXCL12 Leu5 contributes to the stability of the active conformations of TM helices in CXCR4 and ACKR3, we analyzed conformational changes of residues in the TM bundles of the two receptors in simulations with WT CXCL12 and the [P2G]CXCL12. In CXCR4:CXCL12 simulations, CXCL12 Leu5 engages an intramolecular hydrophobic spine in the receptor TM core domain involving Leu41^1.35^, Tyr45^1.39^, and Phe292^7.43^, with the latter controlling the conformation of the intracellular half of TM7 (**Fig. 4A**). In replicate simulations 2 and 3, these residues and their interactions, facilitated by CXCL12 Leu5, maintain CXCR4 in the active state including the “closed” conformation of TM7 (**Fig. 4B**). In CXCR4 simulations with [P2G]CXCL12, the hydrophobic spine collapses due to increased Leu5 dynamics and the loss of contact with Leu41^1.35^; this leads to increased mobility of Tyr45^1.39^ and the loss of its contact with Phe292^7.43^. The intracellular half of TM7 helix rotates counterclockwise (when viewed from the extracellular side) away from TM6 and shifts outwards, breaking the active state of CXCR4 (**Fig. 4A**). These conformational changes were observed in 3/3 simulations of the CXCR4:[P2G]CXCL12 system consistent with the antagonist nature of the CXCL12 variant (**Fig. 4B**). It was also observed in simulation 1 of the CXCR4:CXCL12 system, which showed a transition to an inactive state (**Fig. 2**), paralleling the 31% occupancy of the inactive state by this receptor as observed in smFRET studies (33).

**Figure 4.**
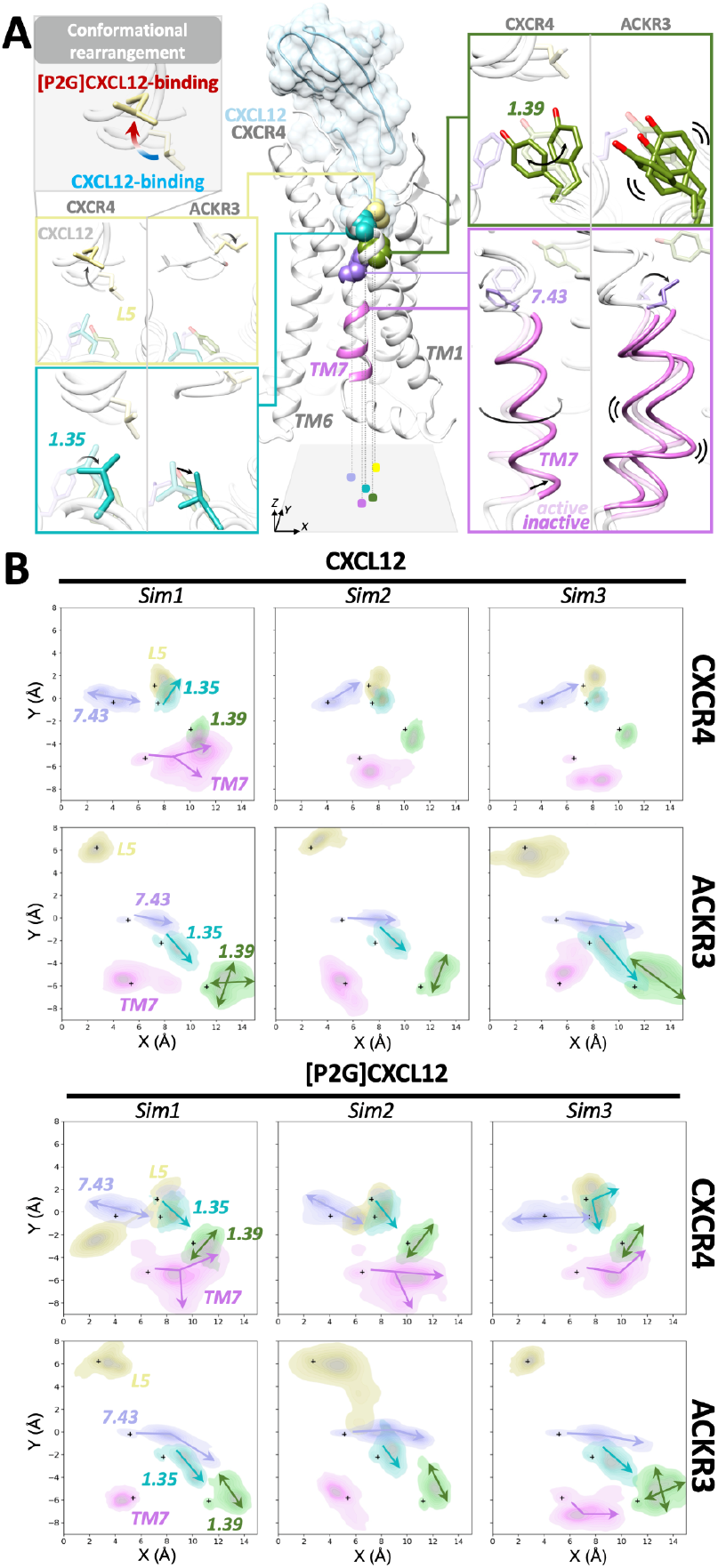
Leu_1.35_-Tyr_1.39_-Phe_7.43_ hydrophobic spine and associated TM7 conformational dynamics in CXCR4 and ACKR3. (**A**) A Leu^1.35^-Tyr^1.39^-Phe^7.43^ network initiated by CXCL12 Leu5 is involved in controlling TM7 conformational changes in CXCR4. This network does not form in ACKR3 leading to a persistently “closed” TM7. Ligand-dependent conformational changes in the hydrophobic spine and TM7 when comparing WT CXCL12-bound (transparent) and [P2G]CXCL12-bound (solid). Inter- and intramolecular hydrophobic spine contacts in CXCR4 and ACKR3 are detailed in the insets. (**B**) Conformational dynamics (n=3) of Leu^1.35^, Tyr^1.39^ and Phe/Leu^7.43^ illustrated with their coordinate distributions within the xy-plane. ‘+’ indicates original conformations.

In ACKR3, Leu47^1.35^ remained highly dynamic regardless of the CXCL12 variant (WT or P2G) due to the lack of an interaction with CXCL12 Leu5. This translated into elevated dynamics of Tyr51^1.39^, allowing the non-aromatic Leu305^7.43^ in ACKR3 to rotate in the opposite direction, relative to Phe292^7.43^ in CXCR4, and leaving TM7 in a persistently closed and active-like conformation (**Fig. 4A-B**). These results suggest that ACKR3 can maintain TM7 in an active state independent of the Leu5-mediated activation network, and independent of the P2G mutation of CXCL12 that produces CXCR4 antagonism.

### ACKR3 lacks the inter-TM aromatic lock required for CXCL12-specific TM6 control

Continuing from the intramolecular hydrophobic spine described in the previous section, in simulations 2 and 3, CXCL12-bound CXCR4 forms a TM1-TM7-TM2-TM3 lock through hydrophobic interaction of aromatic residues Tyr45^1.39^-Phe292^7.43^-Phe87^2.53^-Tyr116^3.32^, which can further regulate TM6 through Leu120^3.36^ and Trp252^6.48^ to maintain its active state (**Fig. 5A**). However, in the CXCR4:CXCL12 simulation 1, which was classified as “inactive” (**Fig. 2**), close contacts of Tyr45^1.39^-Phe292^7.43^ and Phe87^2.53^-Tyr116^3.32^ are broken between either or both pairs (**Fig. 5B**). Similarly, in all [P2G]CXCL12-bound CXCR4 simulations (inactive state), the conformational changes and loss of contacts between Tyr45^1.39^ and Phe292^7.43^ (**Fig. 4**) cause Phe87^2.53^ to move away from Tyr116^3.32^ (**Fig. 5C**).

**Figure 5.**
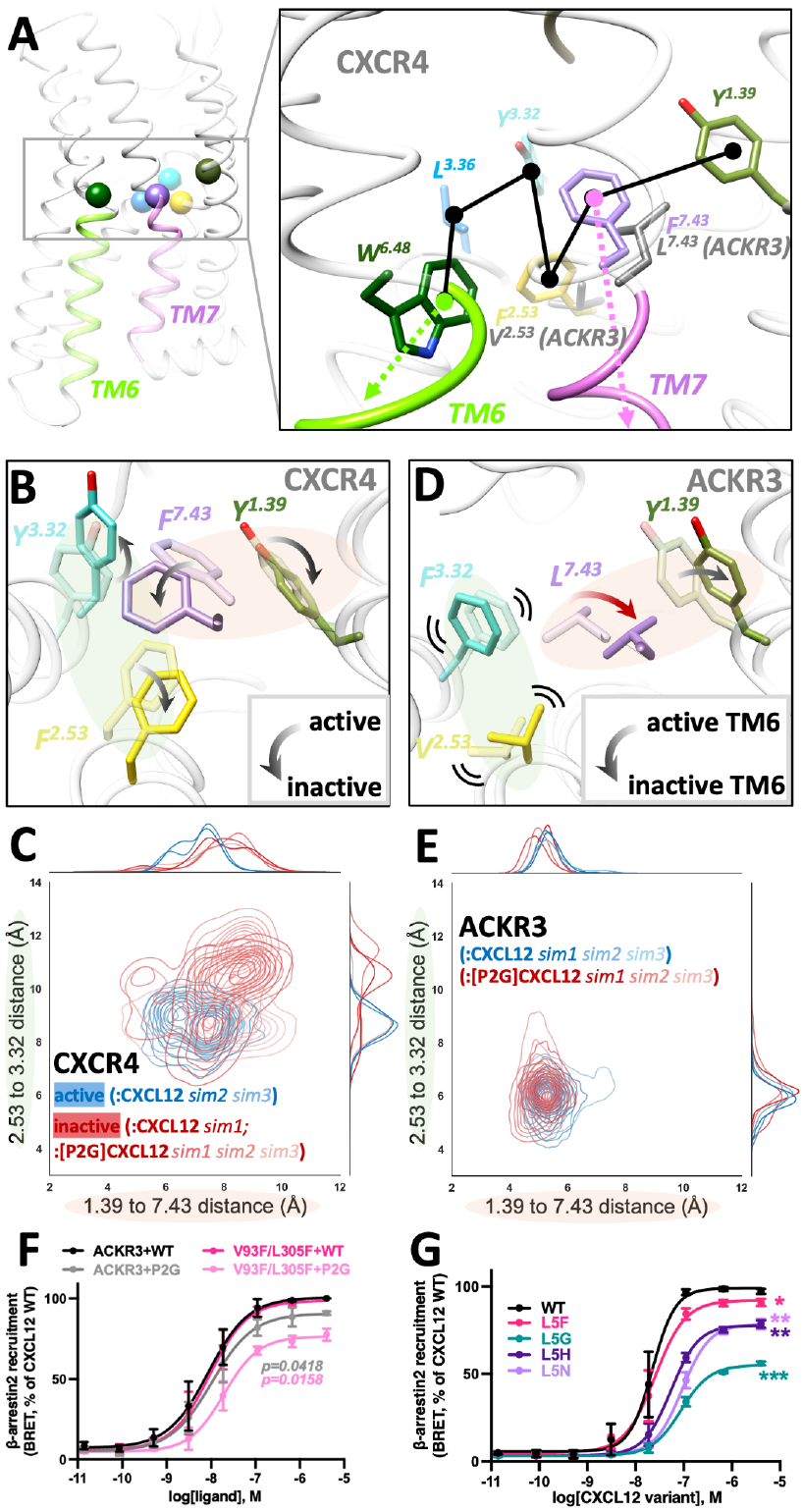
An inter-TM aromatic lock governs TM6 control in CXCR4 but is absent in ACKR3. (**A**)Tyr45^1.39^-Phe292^7.43^-Phe87^2.53^-Tyr116^3.32^ aromatic network involved in TM6 conformational control in CXCR4, which is disrupted by Val93^2.53^ and Leu305^7.43^ substitutions in ACKR3. (**B-E**) Conformational changes and pairwise distances between residues 1.39-7.43 and 2.53-3.32 during simulations. Representative side-chain conformations (**B**) and pairwise distances (**C**) are shown for CXCR4 in active and inactive states. Analogous residue conformations (**D**) and their pairwise distances (**E**) are shown for ACKR3 sampling “active-like” and “inactive-like” TM6 conformations. (**F-G**) BRET measurements in HEK293T cells co-expressing ACKR3-RlucII or ACKR3(V93F/L305F)-RlucII with GFP10-β-arrestin2 and stimulated with varying concentrations of WT CXCL12, [P2G]CXCL12 (**F**) or CXCL12 Leu5 mutants (**G**). Responses were expressed as % maximal response to WT CXCL12 (100%) within each experiment and are presented as mean ± SEM from three independent experiments. Statistical significance was assessed using a one-sample t-test comparing each mutant to the normalized WT response (100%).

Comparative sequence analyses demonstrated that these four aromatic residues are highly conserved in canonical chemokine receptors (**Fig. S14**). However, ACKR3 features non-aromatic residues not only in position 7.43 (Phe292^7.43^ in CXCR4, Leu305^7.43^ in ACKR3), but also in position 2.53 (Phe87^2.53^ in CXCR4, Val93^2.53^ in ACKR3). The loss of these aromatics disrupts the Tyr45^1.39^-Phe292^7.43^-Phe87^2.53^-Tyr116^3.32^-Leu120^3.36^-Trp252^6.48^ network needed for the CXCR4-like TM6 activation (**Fig. 5A**). To assess whether these residues contribute to TM6 dynamics in ACKR3, we compared the conformational changes of Tyr51^1.39^, Leu305^7.43^, Val93^2.53^ and Phe124^3.32^ between an “active-like” TM6 conformation (with a more stable TM6, simulation 1 of ACKR3:[P2G]CXCL12) and an “inactive-like” TM6 conformation (with dynamic TM6, simulation 2 of ACKR3:CXCL12). In the latter, fluctuations of Tyr51^1.39^ promote the movement of Leu305^7.43^ in the same direction as Tyr51^1.39^ via an alkyl–π interaction. However, the Leu305^7.43^ motion fails to propagate to Val93^2.53^ due to its shorter, non-aromatic side chain, leaving Phe124^3.32^ unresponsive to these conformational changes (**Fig. 5D**). The Tyr51^1.39^-Leu305^7.43^ and Val93^2.53^-Phe124^3.32^ contacts remain constant over the course of all ACKR3 simulations with either WT CXCL12 or [P2G]CXCL12, as shown by their stable distances (**Fig. 5E**). These results suggest that the broken aromatic lock impairs conformational coupling between the CXCL12 N-terminus in the orthosteric pocket and the TM6 motions in ACKR3, contributing to TM6 instability.

To verify the roles of Val93^2.53^ and Leu305^7.43^ in the atypical pharmacology of ACKR3, we mutated these residues to their CXCR4 counterparts to generate a double Val^2.53^Phe/Leu^7.43^Phe mutant, and tested its responses to CXCL12 and [P2G]CXCL12 in the β-arrestin recruitment assay. The mutant showed a similar response to WT CXCL12 but significantly lower potency and efficacy of the response to [P2G]CXCL12, suggesting a switch in [P2G]CXCL12 pharmacology to partial agonism (**Fig. 5F, Table S1**). Although Leu5 of CXCL12 plays a limited role in ACKR3 activation, the ACKR3 Val^2.53^Phe/Leu^7.43^Phe mutant also exhibited decreased efficacy and potency in responses to most Leu5 mutants (**Fig. 5G, Table S1**).

WT ACKR3 populates a constitutively active basal state that readily engages β-arrestin (17,18) and undergoes constitutive internalization (19). Interestingly, in addition to altered ligand responsiveness, the Val^2.53^Phe/Leu^7.43^Phe mutant showed higher surface expression and lower basal β-arrestin recruitment than WT ACKR3 (**Fig. S15**). The mutations thus appear to be more conformationally constrained, and shift the receptor toward a more stable inactive-like ensemble, reducing its basal activity.

Together, these results indicate that the aliphatic-to-aromatic mutations in the hydrophobic core of ACKR3 partially restored the canonical conformational coupling between its orthosteric pocket and the intracellular part of its TM bundle, resulting in more CXCR4-like behavior with higher ligand selectivity and lower constitutive activity. However, the Val^2.53^Phe/Leu^7.43^Phe mutant did not exhibit detectable G protein activation under basal or CXCL12-stimulated conditions (**Fig. S16**), indicating that restoration of the aromatic lock is not sufficient to confer G protein coupling to ACKR3.

## Discussion

CXCR4 and ACKR3 share a conserved class A 7TM scaffold and canonical signaling motifs yet exhibit fundamentally distinct activation mechanisms and downstream activities. Although prior structural and biophysical studies have shown that ACKR3 possesses a more dynamic intracellular conformational landscape (33), the molecular basis underlying its divergent functions, including its exceptional susceptibility to activation and its biased intracellular effector engagement has remained unclear. Here, we demonstrate that differences in the non-conserved extracellular chemokine-binding or intracellular effector-binding interfaces are insufficient to explain the atypical pharmacology of ACKR3. Instead, our results indicate that differences in intracellular TM6 and TM7 conformational dynamics, governed by receptor core packing, contribute to its propensity for activation and likely to its inability to couple to G proteins.

Our MD simulations show that CXCL12 binding to CXCR4 stabilizes a coordinated active-state geometry involving TM5, TM6, and TM7, whereas the antagonist [P2G]CXCL12 destabilizes this configuration by opening TM7 and increasing conformational fluctuations in TM6. Thus, activation of CXCR4 involves ligand-dependent stabilization of a well-defined conformational landscape rather than TM6 displacement alone. This agrees with earlier smFRET observations showing that despite its antagonist pharmacology (21), [P2G]CXCL12 shifts CXCR4 toward an active-like TM4-TM6 conformation (33). By contrast with CXCR4, ACKR3 exhibits persistent TM5 and TM6 dynamics regardless of whether CXCL12 or [P2G]CXCL12 is bound, consistent with prior smFRET observations that ACKR3 samples multiple TM6 conformational states (33) (**Fig. S17**). As a result, ACKR3 fails to maintain a canonical fully-outward TM6 conformation in contrast to that of activated CXCR4 and in agreement with a recent HDX-MD study (62). However, it shows a relatively stable active-like positioning of TM7 in response to both ligands, suggesting that the enhanced conformational heterogeneity of ACKR3 arises not simply from increased TM6 mobility, but from weakened coordination between TM6 and TM7.

To identify the structural basis for the weakened TM6-TM7 coordination in ACKR3 compared to CXCR4, we examined the TM core networks that stabilize the active conformations. Activation of CXCR4 appears to require coordinated stabilization of a conserved TM1-TM7-TM2-TM3 aromatic lock at BW positions 1.39, 7.43, 2.53, and 3.32, which further govern the conformation of TM6 (**Fig. 6A**). This lock constrains the receptor into a well-defined active conformation upon CXCL12 binding, including a rotated and closed TM7 configuration and a stabilized TM6. Even subtle perturbations in the upstream ligand-receptor interaction network, such as changes in CXCL12 N-terminal residues Lys1 or Pro2 (21,57), or mutations to Trp94^2.60^, Asp97^2.63^, Tyr116^3.32^, Asp171^4.60^, and Asp187^ECL2^ (20,23,36,37,50,51,53–56) can disrupt this aromatic lock, and hence CXCR4 activation, without detrimental effects on CXCR4-CXCL12 binding. This explains why [P2G]CXCL12 is a pharmacological antagonist: mutation of Pro2 leads to loss of coordinated control between TM6 and TM7, resulting in an unrotated and more open TM7 conformation and increased TM6 fluctuations, but preserves binding. In our study, we also demonstrate that this holds for mutations in CXCL12 Leu5, a residue without previously known roles in CXCR4 activation. In ACKR3, this aromatic lock is disrupted due to the absence of aromatic residues at positions 7.43 and 2.53 (where they are replaced by branched aliphatics Leu305^7.43^ and Val93^2.53^). Aliphatic residues exhibit more permissive packing interactions compared with the directional π–π stacking between the aromatic side chains (63,64) (Phe292^7.43^ and Phe87^2.53^ in CXCR4). These substitutions result in a more conformationally flexible TM core in ACKR3, thereby reducing the energetic barrier between conformational states and making ACKR3 activation-prone. This atypical behavior is reflected by a persistently unrotated and opened TM7 and unconstrained TM6 dynamics, regardless of mutations in the CXCL12 N-terminus. Similar mechanisms may also enable ACKR3 activation by other ligands of varying sequence and size (16,29–32). Together, these results support a model in which CXCR4 is activated by a structurally constrained network of interactions between CXCL12 and the TM core, whereas the flexible TM core of ACKR3 enables activation by diverse ligands through a less specific distortion-based mechanism (**Fig. 6B**), as we previously suggested (33,34). Consistent with this, a recent study of ACKR3 bound to inverse agonist-, antagonist-, partial agonist-, and agonist-associated nanobodies revealed that increasing receptor activity correlates with progressive opening of the extracellular orthosteric pocket (18).

**Figure 6.**
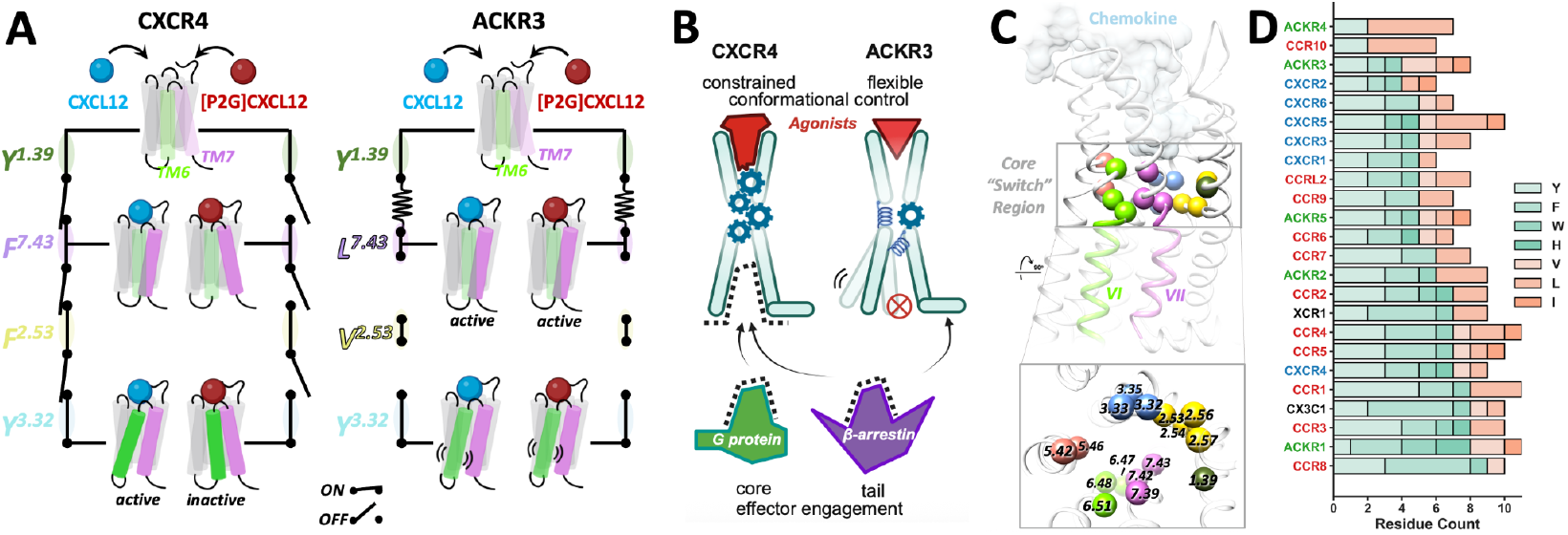
Core packing tunes TM dynamics and receptor behavior. (**A**) Conceptual model illustrating how non-conserved aromatic lock residues influence TM6/TM7 conformational dynamics in ACKR3. (**B**) Proposed model showing that CXCR4 adopts a constrained TM core which supports ligand-dependent and tightly controlled effector binding, whereas ACKR3 exhibits a loosened core that contributes to its activation prone nature and possibly its impaired effector coupling. (**C**) Structural model highlighting the proposed “core-switch region” of class A GPCRs and the corresponding BW positions that may form “aromatic locks”. (**D**) Stacked bar plot showing the frequency of aromatic-like (Y, F, W, H) and branched aliphatic hydrophobic (V, L, I) residues at the BW positions indicated in (B) across chemokine receptors. Receptors are ranked by aromatic-like residue count.

The distinct cytoplasmic TM dynamic landscape may also be responsible for the differential intracellular effector coupling of CXCR4 and ACKR3 (**Fig. 6B**). Our analysis indicates that TM4-TM6 separation, while associated with active conformations, is not sufficient on its own to support canonical G protein coupling. Consistent with this, a recent systematic MD study indicates that both the location (spatial) and duration (temporal) of intermolecular contacts at the GPCR:Gα interface determine how GPCRs engage G proteins (65). We show that ACKR3 fails to occupy a stable outward-facing conformation of TM6 and, consequently, lacks a well-defined and persistent intracellular effector-coupling interface, explaining its inability to activate G proteins. The lack of intracellular structural organization may also explain why ACKR3 interacts with β-arrestin only with its C-tail but not the cytoplasmic TM cavity (66); this is unlike canonical GPCRs (rhodopsin, neurotensin receptor 1, 5-HT2B, β1 adrenoceptor, M2 muscarinic receptor, and vasopressin V2 receptor) that form a fully engaged “core-and-tail” complex (67–73). ACKR3 tail interaction with β-arrestin does, however, depend on its phosphorylation by GRK5 or GRK2 (9,19,35). The abrogation of GRK2/5-mediated C-tail phosphorylation by CID24, a Fab that blocks the intracellular TM cavity of ACKR3 (66), suggests that in contrast to G proteins and β-arrestins, GRKs may couple to ACKR3 through its cytoplasmic TM pocket, similar to rhodopsin with GRK1 (74), neurotensin receptor 1 with GRK2 (75), and β2 adrenoceptor with GRK5 (76). Whatever the precise binding mode, GRK-mediated phosphorylation enables the C-tail-only ACKR3 association with β-arrestin while the cytoplasmic pocket dynamics may prevent G protein coupling, resulting in the complete bias of ACKR3 towards β-arrestin.

The current study highlights the critical role of TM core packing in modulating chemokine receptor responsiveness to ligands, with aromatic residues contributing to a more conformationally constrained core and aliphatic residues favoring increased flexibility, consistent with the idea that less geometrically constrained hydrophobic packing can facilitate conformational dynamics (77–79). These findings may also have broader implications for chemokine receptor activation mechanisms and cytoplasmic effector bias. To explore whether variation in core packing composition is associated with receptor behavior across the chemokine receptor family, we quantified the distribution of aromatic (Phe, Tyr, Trp, His, planar ring-containing and capable of π-stacking) and branched aliphatic (Val, Leu, Ile, less directional packing) residues within a defined TM “core switch region” (**Fig. 6C-D**). ACKR3 exhibits reduced aromatic content and increased aliphatic composition in this region compared to most chemokine receptors, consistent with its enhanced conformational flexibility. ACKR4, another β-arrestin-biased atypical chemokine receptor with scavenging function (80), shows an even greater shift toward aliphatic residues, suggesting that this atypical receptor may share a common structural basis for its unconventional effector coupling. In addition, CXCR2, a canonical receptor and one of the most promiscuous members of the CXC subfamily (81), can be activated by chemokines with varying lengths and compositions of N-termini, with the most potent agonists featuring only a shallow insertion of the N-terminus into the TM binding pocket. Rather than forming a well-defined and ‘locked’ network, the N-termini of CXCR2-activating chemokines (CXCL1, CXCL2, CXCL3, CXCL5, CXCL6, and CXCL8) adopt extended conformations and primarily engage the extracellular vestibule (82). CXCR2 exhibits reduced aromatic content in its TM core, which may contribute to a more permissive activation, similar to ACKR3. On the other hand, with the exception of CCR10, receptors within the CCR subfamily exhibit higher aromatic content within the TM core, compared to many CXCRs, yet display a wide range of ligand promiscuity (e.g., CCR1 (83) and CCR3 (84)). Moreover, several atypical chemokine receptors, including ACKR1, ACKR2, and ACKR5, also maintain relatively aromatic-rich TM cores yet exhibit non-canonical effector coupling (85–87). This diversity suggests that core packing composition alone does not fully account for chemokine responsiveness and effector engagement. Specific aromatic residue positioning within the TM bundle and additional structural determinants, including conserved TM motifs (e.g., disrupted or non-canonical DRY and NPxxY motifs in ACKR1 and ACKR2 (88)), ECL architecture (58), chemokine N-terminal engagement modes (89), orthosteric pocket geometry (90), and intracellular architectures and effector interactions (18,65), can also modulate how chemokine binding is coupled to intracellular conformational changes. This is consistent with our observation that ACKR3 Val^2.53^Phe/Leu^7.43^Phe mutant still does not exhibit detectable G protein activation.

Altogether, the present study demonstrates how dynamics can regulate receptor activation energetics and responsiveness to ligands, and potentially effector interaction bias. These principles may extend to other atypical receptors as well as some dual-function chemokine receptors (e.g., CCR2 (91)), where conformational dynamics could modulate a functional spectrum spanning G protein signaling and atypical scavenging-like behavior.

## Materials and methods

### Reagents and supplies

Commercially available reagents and supplies used in this work are listed in **Table S2**.

### DNA constructs and site-directed mutagenesis

For CXCL12 production, a pMS211-based construct of human CXCL12α (residues Lys22–Lys89) lacking the endogenous N-terminal signal peptide and preceded by an enterokinase recognition site and His8 tag (59,92), was used as the WT construct. For clarity, Lys22 of CXCL12 is renumbered as Lys1 throughout this study. For the G protein dissociation assay, pcDNA3.1 expression constructs encoding ACKR3 or CXCR4 were generated in our lab (42,51). An N-terminal FLAG tag was added to ACKR3 while CXCR4 had an N-terminal HA tag. The tricistronic pIRES-Gβ1-2A-cpVenus-Gγ2-Gαi1-Nluc plasmid was gifted from Professor Daniel Legler from University of Konstanz. For the β-arrestin recruitment assay, the FLAG-ACKR3 or HA-CXCR4 segments were fused with a C-terminal Renilla luciferase II (RlucII, receptor-RlucII) and that of GFP10 fused to human β-arrestin2 (GFP10-β-arrestin2), both in pcDNA3.1 vectors, followed by cloning of all mutants. Site-directed mutagenesis was performed using Q5 Site-Directed Mutagenesis Kit (New England Biolabs, Ipswich, MA). Point mutations were introduced into ACKR3 and CXCL12 plasmids by PCR mutagenesis. All mutations were confirmed by Sanger sequencing.

### Cell culture

HEK293T cells were obtained from the American Type Culture Collection (ATCC) and certified mycoplasma-free. The cells were cultured in sterile tissue culture treated 10 cm dishes in Dulbecco’s Modified Eagle Media (DMEM) with 10% fetal bovine serum (FBS) at 37°C in a humidified incubator with 5% CO_2_. Cells were routinely passaged 1:10 every 2 to 3 days using 0.05% trypsin-EDTA. Functional experiments were restricted to cells with no more than 15 passages.

### Chemokine expression and purification

The chemokine expression and purification protocol was adapted from (59). Plasmids encoding CXCL12 and CXCL12 mutants were transformed into BL21(DE3)pLys competent cells. Cells were grown at 37°C with shaking at 250 rpm to an optical density at 600 nm of 0.6 to 0.7 in Luria-Bertani medium, and protein expression was induced by addition of 1 mM isopropyl β-D-1-thiogalactopyranoside. Following 6 h of induction at 37°C, cells were harvested by centrifugation. Cell pellets were resuspended in lysis buffer (50 mM Tris, 150 mM NaCl [pH 7.5]) with 1 μg/mL DNaseI and lysed by sonication. The samples were centrifuged (12,000g for 15 min), and the insoluble pellets containing chemokine inclusion bodies were collected. The pellets were dissolved in Equilibration buffer (50 mM Tris, 6 M guanidine-HCl, 50 mM NaCl [pH 8.0]), sonicated, and the supernatant (containing chemokine) was collected by centrifugation (12,000g for 15 min). The supernatant was loaded onto Ni-NTA agarose resin (Qiagen, Hilden, Germany), washed with Wash buffer (50 mM MES, 6 M guanidine-HCl, 50 mM NaCl [pH 6.0]), and the chemokine was eluted with Elution buffer (50 mM acetate, 6 M guanidine-HCl, 50 mM NaCl [pH 4.0]). The eluate was brought to pH > 7.0 with NaOH, reduced with 4 mM DTT for 2 h at room temperature, diluted 10-fold into Refolding buffer (50 mM Tris, 500 mM arginine-HCl, 1 mM EDTA, 1 mM glutathione disulfide [pH 7.5]), and incubated for 60 min at room temperature with moderate stirring. Following dialysis of the refolding mixture (20 mM Tris, 50 mM NaCl [pH 8.0]), the dialysis product was cleared of precipitant by centrifugation (8,000g for 15 min) and concentrated to approximately 10 to 20 mL with a 3 kDa cutoff Amicon centrifugal filter unit (MilliporeSigma, Burlington, MA). After concentration, 2 mM CaCl_2_ was added and the His8 tag was cleaved through the addition of 8 to 16 U/mL enterokinase (New England Biolabs, Ipswich, MA) for 2 days at 37°C. After cleavage, the mixture was loaded onto Ni-NTA resin, and tag-free CXCL12 was eluted with Wash buffer. The eluate was loaded onto a reversed-phase C18 HPLC column (Vydac, Columbia, MD; buffer A: 0.1% trifluoroacetic acid; buffer B: 0.1% trifluoroacetic acid; 90% acetonitrile) in 67% buffer A/33% buffer B, and eluted by linearly increasing buffer B from 33% to 45%. The resulting chemokine was lyophilized and stored at −80 °C until use. Lyophilized chemokine was dissolved in sterile distilled water, and the protein concentration was determined by measuring absorbance at 280 nm using a NanoDrop spectrophotometer (Thermo Fisher Scientific, Waltham, MA).

### BRET to determine CXCL12-stimulated G protein dissociation

HEK293T cells were seeded directly into 96-well tissue culture plates (50,000 cells per well) and transfected after 6 h using Mirus TransIT-LT1 transfection reagent (Mirus Bio, Madison, WI). The cells were transfected with 50 ng of pcDNA3.1-receptor per well, along with 50 ng of tricistronic pIRES-Gβ1-2A-cpVenus-Gγ2-Gαi1-Nluc plasmid. For selected receptor mutants exhibiting reduced cell surface expression, including ACKR3 chimeric mutants containing the N-terminus and ECL2 of CXCR4, the amount of receptor plasmid was increased to 100 ng per well to partially compensate for impaired surface expression (**Fig. S3**). Cells were treated with 100 ng/ml PTX (Thermofisher/Invitrogen, Carlsbad, CA) at 24 h post-transfection to measure the basal coupling of Gαi to the GPCRs (40). BRET assays were performed ∼42 h after transfection, using a slight modification of a previously described method (20). Cells were washed once with pre-warmed Tyrode’s buffer (25 mM HEPES, 140 mM NaCl, 2.7 mM KCl, 1 mM CaCl_2_, 12 mM NaHCO_3_, 5.6 mM D-glucose, 0.5 mM MgCl_2_, 0.37 mM NaH_2_PO_4_ [pH 7.4]), then the cell-permeable Nluc substrate, coelenterazine-h (NanoLight Technologies, Pinetop, AZ) was added at a final concentration of 5 μM ∼3 min before BRET measurements. Three baseline BRET measurements were performed approximately 1 min apart and basal BRET was recorded, followed by the addition of 100 nM CXCL12. Subsequent BRET measurements were taken approximately 1 min apart for 20 min. BRET measurements were made on a TECAN Spark Luminometer (TECAN Life Sciences, Männedorf, Switzerland) at 37 °C using default BRET1 settings (blue emission 430-485 nm, red emission 505–590 nm) and an integration time of 0.2 s. The collected luminescence emissions were converted into BRET ratio, baseline-corrected by subtracting the average BRET ratio prior to agonist addition, and normalized as described in the figure legends. Assays were performed in three independent biological replicates, each with 2–3 technical replicates.

### BRET to determine CXCL12-stimulated β-arrestin recruitment to receptors

HEK293T cells were seeded directly into 96-well tissue culture plates (50,000 cells per well) and transfected after 6 h using Mirus TransIT-LT1 transfection reagent (Mirus Bio, Madison, WI). Cells were transfected with the BRET donor (5 ng of receptor-RlucII per well) along with the BRET acceptor (95 ng of GFP10-β-arrestin2 per well). All assays were performed ∼42 h after transfection, with slight modifications from previously described methods (93). Cells were washed once with pre-warmed Tyrode’s buffer (25 mM HEPES, 140 mM NaCl, 2.7 mM KCl, 1 mM CaCl_2_, 12 mM NaHCO_3_, 5.6 mM D-glucose, 0.5 mM MgCl_2_, 0.37 mM NaH_2_PO_4_ [pH 7.4]), then cell-permeable RlucII substrate, Prolume Purple (NanoLight Technologies, Pinetop, AZ) was added at a final concentration of 5 μM ∼2 min before BRET measurements. Two baseline BRET measurements were performed approximately 2.5 min apart, followed by the addition of the indicated concentrations of chemokine. Subsequent BRET measurements were taken approximately 2.5 min apart for 30 min. BRET measurements were made on a TECAN Spark Luminometer at 37 °C using default BRET2 settings (blue emission 360–440 nm, red emission 505–575 nm) and an integration time of 0.1 s. The collected luminescence emissions were converted into BRET ratio, baseline-corrected by subtracting the average BRET ratio prior to agonist addition, and normalized as described in figure legends. Dose–response curves were analyzed using nonlinear regression (four-parameter logistic model) to derive EC50 and Emax values using GraphPad Prism version 10.5.0 (GraphPad Software, San Diego, CA). Assays were performed in three independent biological replicates, each with 2–3 technical replicates.

### Flow cytometry-based surface expression testing

The cell surface expression of WT and mutants of ACKR3 and ACKR3-RlucII was monitored by flow cytometry as described previously (51). HEK293T cells were seeded directly into 96-well tissue culture plates (50,000 cells per well) and transfected after 6 h using Mirus TransIT-LT1 transfection reagent (Mirus Bio, Madison, WI). For ACKR3 chimeric mutants used in G protein dissociation BRET assays, the same amount of receptor plasmid was used for both the flow cytometry and the corresponding BRET experiment. For the ACKR3 V93F/L305F mutant used in β-arrestin recruitment BRET assays, 50 ng of receptor was transfected to enable detection of cell surface expression by flow cytometry, as lower DNA amounts used in BRET assays resulted in signals below the detection threshold. Forty-eight hours after transfection, cells were washed with PBS, lifted with Accutase (STEMCELL Technologies, Vancouver, Canada), and resuspended in cold FACS buffer (PBS, 0.5% BSA). Every 1 × 10^5^ cells were added with 30 ng of APC-conjugated anti-FLAG antibody (clone L5, BioLegend, San Diego, CA) for the detection of the N-terminal FLAG tags in ACKR3 chimeric mutants, or PE-conjugated anti-ACKR3 antibody (clone 11G8, R&D Systems, Minneapolis, MN) to obtain a 30X dilution for the detection of ACKR3 V93F/L305F. Cells were stained on ice in the dark for 30 min, then washed three times and resuspended to ∼5 × 10^5^ cells/ml with the FACS buffer. All staining and washing procedures were performed on ice and with ice-cold buffers to avoid receptor internalization. Flow cytometric analysis of the antibody-stained cells was performed with a GUAVA benchtop flow cytometer (EMD Millipore, Burlington, MA). Flow cytometry data was analyzed with FlowJo version 10.8.1 (FlowJo, Ashland, OR), and relative fluorescence geometric mean (RFGM) values were normalized to those of WT ACKR3 (100%) after subtraction of the RFGM obtained for cells transfected with same amount of pcDNA and stained with the same antibody.

### Preparation of initial receptor:chemokine models for MD simulations

Structural models of ACKR3 and CXCR4 complexes with WT CXCL12 were constructed by AlphaFold2 Multimer v2.3.2 (41,94) locally installed on the UCSD Triton Shared Computing Cluster (TSCC). An ensemble of 25 models (5 random seeds with 5 models per seed) was built for each of the complexes using default settings. Models were analyzed as in (95) taking into account the interaction prediction confidence (chemokine N-terminus pLDDT and inter-chain PAE (41,43)) and interaction favorability (RTCNN (96,97)). The top-scoring model was selected as a starting point for the simulation.

To avoid unwanted interactions between receptor terminal sequences across periodic boundaries that we observed in preliminary simulations (data not shown), receptor termini were truncated using ICM Pro version 3.9-4a (Molsoft LLC, San Diego, CA) (98) For ACKR3, residues Met1-Ile17 of ACKR3 were removed, creating a construct that experimentally retains full receptor activity (42,59) (**Fig. S2, Fig. S12**). For CXCR4, Met1 and Glu2 were removed to prevent dissociation of the CXCR4 N-terminus from CXCL12 caused by electrostatic repulsion between Glu2 and negatively charged Glu179, Asp181, and Asp182 in ECL2 (**Fig. S2, Fig. S12**). In addition, flexible sequences following H8 were removed (residues Ala336-Lys362 in ACKR3 and residues Arg322-Ser352 in CXCR4). ICM Pro was also used to introduce the P2G mutation in CXCL12 for the [P2G]CXCL12-bound models. Each system was subjected to energy minimization with restraints in ICM, to remove minor steric clashes.

### MD simulation system setup

The above models of ACKR3 and CXCR4:CXCL12 complexes of ACKR3 and CXCR4 with CXCL12 and [P2G]CXCL12 were used as starting structures for MD system preparation. Prior to system assembly, the complexes were positioned relative to the membrane normal by superimposing the receptor TM backbone onto a precomputed and pre-oriented collection of GPCR structures with cholesterol molecules from the PDB, to ensure a consistent and physiologically relevant in-membrane orientation. Membrane insertion, solvation, and neutralization were performed in CHARMM-GUI Membrane Builder (99). ACKR3 and CXCR4 were capped with an N-terminal acetyl group and a C-terminal N-methylamide. All chemokines were C-terminally capped with an N-methylamide, while their N-termini were kept intact and charged. The Cys34-Cys287 disulfide bond in ACKR3 (present in the AF2 model) was removed in accordance with the cryo-EM structure (PDB 7SK3) (34). Residues were protonated at pH 7.0. His79^2.45^ of CXCR4 was manually tautomerized as HSE to stabilize the TM2-TM4 interaction via a His79^2.45^-Trp161^4.50^ hydrogen bond, consistent with previous structural models (PDBs 8U4N, 8U4R, and 8K3Z). Protein complexes were embedded in a POPC bilayer (∼75 lipids per leaflet) with a simulation box of approximately 80 × 80 × 135 Å^3^ in the x, y, and z dimensions. For CXCR4 systems, the box was extended to ∼150 Å along the z-axis to prevent unwanted interactions between the flexible N-terminus and periodic images. Systems were solvated with TIP3P water and neutralized with Na^+^ and Cl^+^ ions to a final salt concentration of 150 mM. The *viparr* tool (D.E. Shaw Research, New York, NY) (100) was used to parameterize proteins, lipids, water, and ions in the CHARMM36m force field (101), and to apply constraints on covalent bonds involving hydrogen atoms using the SHAKE algorithm (102).

### System relaxation

All-atom MD relaxations and simulations were performed using Desmond (D.E. Shaw Research, New York, NY) (103) on the UCSD TSCC. The systems were relaxed through a multistep equilibration protocol, initially developed by D. Lupyan and Schrödinger scientists and supplied with the academic distribution of Desmond (103). Initial relaxation employed low-temperature Brownian dynamics in the NVT ensemble (T = 10 K), with positional restraints applied to all solute heavy atoms. The temperature was gradually increased stepwise to 100 K under NVT conditions with a Nosé–Hoover thermostat, while restraints were maintained on the protein and partially on the membrane along the z-axis. Subsequent equilibration involved gradual heating to 310 K using NPT and then NPgT ensembles (P = 1 bar) with MTK or Langevin methods, during which Gaussian forces were applied to solvent molecules to facilitate water relaxation. The last two equilibration phases over ten nanoseconds involved exponentially decreased restraints on proteins. Positional restraints of proteins were gradually released in a sequential manner: side chains first, followed by non-TM backbone atoms including H8, and finally TM helices, allowing proper H8-membrane interaction and progressive relaxation of the protein in the membrane. The cumulative equilibration time across all stages was ∼20 ns.

### Production MD simulation

Following system relaxation, production MD simulations were performed under the NPgT ensemble at 310 K and 1 bar using a Langevin thermostat (τ = 1.0 ps) and barostat (τ = 2.0 ps). No positional restraints were applied. Initial velocities were randomly assigned according to a Maxwell–Boltzmann distribution at 310 K (seed = 2007). Simulations were conducted for 10 μs with an integration timestep of 2 fs for bonded and short-range non-bonded interactions, and 6 fs for long-range interactions. Short-range interactions were treated with a 9 Å cutoff. Long-range electrostatic interactions were treated using the smooth particle mesh Ewald method (104). Coordinates and velocities were saved at 50 ps intervals in the DTR format. Periodic boundary conditions were applied, and simulation box dimensions were recorded every 1.2 ps. Checkpoints were written every 240 ps to allow restart if necessary. Independent simulations in replicates of three for each system were initiated using different randomized initial velocities.

### Trajectory processing and analysis

Trajectories were wrapped and centered on the receptor TM backbone using the *molfile* and *pfx* modules of the DESRES *msys* Python library version 1.7.359 (100). Receptor orientation relative to the membrane (**Fig. S5**) was quantified by measuring the angle between the principal axis of the TM bundle and the z-axis of the coordinate system. Prior to receptor RMSD and conformational analyses, the trajectories were aligned on the receptor TM backbone using *msys*.*molfile* and *msys*.*pfx. Msys*.*molfile* was also used to strip solvents and ions and to sparsely subsample the trajectory (∼5 ns/frame unless otherwise stated) for efficient downstream analyses. System stability was visually assessed in VMD version 1.9.2 (105) and quantified by measuring the RMSD of protein backbone atoms (**Figs. S6** and **S7**) in Python *MDAnalysis* library version 2.7.0 (106,107). The integrity and dynamics of the POPC bilayer (bilayer thickness, area per lipid, and lipid lateral diffusion, **Figs. S8, S9**, and **S10**, respectively) were analyzed using the *lipyphilic* Python library version 0.11.0 (108). RMSD, pairwise inter- and intramolecular distances, and coordinates shown in other figures were measured using *MDAnalysis* and plotted in Python using *matplotlib* version 3.7.1 and *seaborn* version 0.12.2.

### Visualization and figure preparation

Structural models and representative trajectory snapshots were visualized and rendered using UCSF Chimera version 1.15 (109). Functional assay data were analyzed using GraphPad Prism version 10.5.0. Schematic diagrams of molecular interactions and proposed models were prepared with BioRender (https://biorender.com).

### Statistical analyses

Data analysis was performed using GraphPad Prism version 10.5.0. Dose–response curves were fitted using nonlinear regression (four-parameter logistic model). Statistical differences in EC50 values between WT and mutants were assessed using an extra sum-of-squares F test. For datasets normalized to WT (set to 100%), including β-arrestin recruitment, basal luminescence, and selected surface expression measurements, statistical significance was assessed using one-sample t-tests comparing each condition to the WT reference value. Unless otherwise stated, data are presented as mean ± SEM from three independent experiments. Significance levels are indicated as not significant (ns), *p* < 0.05 (*), *p* < 0.01 (**), *p* < 0.001 (***), and *p* < 0.0001 (****).

## Abbreviations

CXCR: C-X-C chemokine receptor
CXCL12: C-X-C chemokine ligand 12
ACKR: Atypical chemokine receptor
MD: Molecular dynamic
WT: Wild type
TM: Transmembrane
BRET: Bioluminescence resonance energy transfer
GPCR: G protein coupled receptor
GRK: G protein-coupled receptor kinase
Cryo-EM: Cryogenic electron microscopy
smFRET: Single-molecule fluorescence resonance energy transfer
ECL: Extracellular loop
ICL: Intracellular loop
H8: Helix 8
PDB: Protein data bank
PTX: Pertussis toxin
SMO: Smoothened receptor
AF2: AlphaFold2
RMSD: Root mean square deviation
AF3: AlphaFold3
POPC: 1-palmitoyl-2-oleoyl-sn-glycero-3-phosphocholine
CRS: Chemokine recognition site
COM: Center-of-mass
SEM: Standard error of the mean
BW: Ballesteros-Weinstein
CCR: C-C chemokine receptor
HA: Hemagglutinin
RLuc: Renilla Luciferase
PCR: Polymerase chain reaction
HEK293T: Human Embryonic Kidney 293T cells
ATCC: American Type Culture Collection
DMEM: Dulbecco’s Modified Eagle Media
FBS: Fetal bovine serum
DTT: Dithiothreitol
EDTA: Ethylenediaminetetraacetic acid
HPLC: High-performance liquid chromatography
PE: Phycoerythrin
APC: Allophycocyanin
RFGM: Relative fluorescence geometric mean
TSCC: Triton Shared Computing Cluster
pLDDT: Predicted local distance difference test
RTCNN: Radial and topological convolutional neural network score

## Acknowledgements

We thank Drs. Knut Teigen and Aage Skjevik (University of Bergen, Norway) for help with initial CXCR4 simulations and Dr. Hishara K. Gallage Dona (UC San Diego / Univ. of Utah) for piloting the *lipyphilic* python library. We also thank members of the Kufareva and Handel labs for valuable and insightful discussions. We acknowledge the San Diego Supercomputer Center (SDSC) for providing computational resources through the Triton Shared Computing Cluster (TSCC). We also thank the Pittsburgh Supercomputing Center (PSC), D. E. Shaw Research (DESRES), and the Anton-2/3 (110,111) training workshop organizers (supported by NIH grants R01GM116961 and R24GM154042) for guidance on simulation system preparation and force-field assignment workflows.

This work was supported by NIH R01CA254402 to T.M.H., R01AI161880 to T.M.H. and I.K., R21AI149369 and R21AI156662 to I.K., the UC Office of the President Cancer Research Coordinating Committee (CRCC) seed grant C26CR10041 to I.K., and the CSD/UCSF Cancer Cell Mapping Initiative Pilot Grant (under NIH U54 CA274502) to I.K. K.W. is supported by an American Heart Association Postdoctoral Fellowship (26POST1563101).

## Author contributions

I.K. and T.M.H. conceived the study. I.K., T.M.H. and K.W. designed the research. K.W. and T.N. cloned receptor mutants and cloned, expressed, and purified WT and mutant CXCL12. K.W., T.N., E.K., S.R. and C.T.S. performed cell-based experiments. R.C. and I.K. performed AF2 modeling and initial MD optimization. T.N. piloted the analysis of initial CXCR4 simulations. K.W. performed MD optimization and simulations used in the current study. K.W. analyzed experimental data and MD trajectories. K.W. and I.K. contributed MD analytical tools. K.W. prepared the figures and visualizations. K.W., I.K. and T.M.H. wrote the manuscript. I.K. and T.M.H. supervised the study. All authors reviewed and approved the final manuscript.

