## Supplemental Information for "Elevated conformational dynamics makes ACKR3 activation-prone and G protein-incompetent"

A

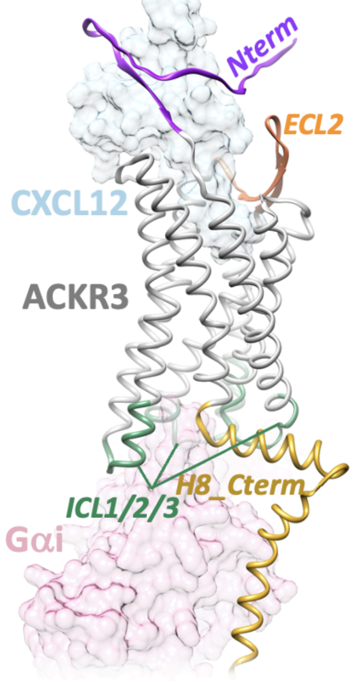


B

```
CXCR4     MEGISIYTSDNYTEEMGSGDYDSMKEPCFREENANFN-------------KIFLPTIYSI 47
ACKR3     -------MDLHLFDYSEPGNFSDISWPCNSSDCIVVDTVMCPNMPNKSVLLYTLSFIYIF 53
                 . :  :    *::..:. **  .:    .:              *  ** :
                          Nterm                                 TM1

CXCR4     IFLTGIVGNGLVILVMGYQKKLRSMTDKYRLHLSVADLLFVITLPFWAVDAV--ANWYFG 105
ACKR3     IFVIGMIANSVVVWVNIQAKTTGYDTHCYILNLAIADLWVVLTIPVWVVSLVQHNQWPMG 113
          **: *::.*.:*: *    *.     *. * *:*::*** .*:*:*.*.*. *   :* :*
                            ICL1                TM2

CXCR4     NFLCKAVHVIYTVNLYSSVLILAFISLDRYLAIVHATNSQRPRKLLAEKVVYVGVWIPAL 165
ACKR3     ELTCKVTHLIFSINLFGSIFFLTCMSVDRYLSITYFTNTPSSRKKMVRRVVCILVWLLAF 173
          :: **..*:*:::**:.*:::*: :*:****:*.: **:    ** :..:** : **: *:
                TM3                                 ICL2                TM4

CXCR4     LLTIPDFIFAN--VSEADDRYICDRFYPND---LWVVVFQFQHIMVGLILPGIVILSCYC 220
ACKR3     CVSLPDTYYLKTVTSASNNETYCRSFYPEHSIKEWLIGMELVSVVLGFAVPFSIIAVFYF 233
           :::**  : :  .* :::.  *  ***:.     *:: :::   :::*: :*  :*   *
                       ECL2                                 TM5

CXCR4     IIISKLSHSKGHQKRKALKTTVILILAFFACWLPYYIGISIDSFILLEIIKQGCEFENTV 280
ACKR3     LLARAISASSDQEKHSSRKIIFSYVVVFLVCWLPYHVAVLLDIFSILHYIPFTCRLEHAL 293
          ::   :* *..::*:.: *  .  ::.*:.*****::.: :* * :*. *   *.:*:::
                 ICL3                    TM6

CXCR4     HKWISITEALAFFHCCLNPILYAFLGAKFKTSAQHALTSV-SRGSSLKILSKGKRGGHSS 339
ACKR3     FTALHVTQCLSLVHCCVNPVLYSFINRNYRYELMKAFIFKYSAKTGLTKLIDASRVSE-- 351
          .. : :*:.*::.***:**:**:*:. ::: .  :*:     *  :.*. *  ...* ..
          TM7                       ⊔              H8_Cterm
                             TM7-H8 hinge
CXCR4     VSTESESSSFHSS-- 352
ACKR3     ----TEYSALEQSTK 362
              :* *:::..*
```

**Figure S1. Amino acid sequence alignment of CXCR4 and ACKR3.** Amino acid sequences were aligned using Clustal Omega implemented in the EMBL-EBI multiple sequence alignment web server. Transmembrane (TM) regions are shown in bold. ACKR3 residues substituted with the corresponding residues from CXCR4 or CXCR2 used for G protein dissociation assay are colored and underlined. "*", ":", and "." indicate identical, highly conserved, and weakly conserved residues, respectively.

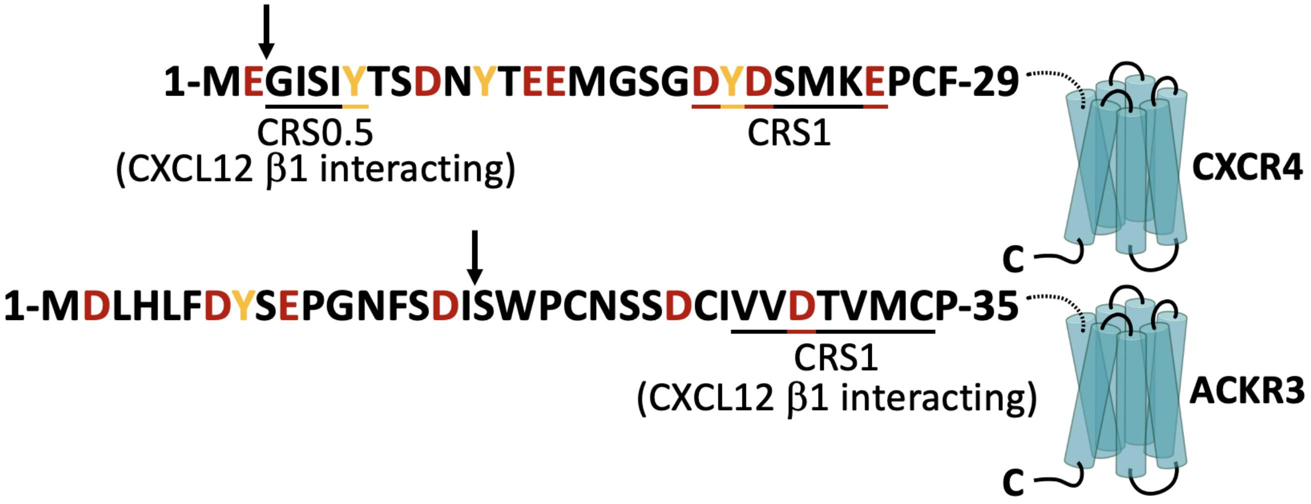


**Figure S2. N-terminal sequences of CXCR4 and ACKR3.** The ACKR3 N-terminus is longer and displays an altered distribution of acidic residues (red), tyrosines (yellow) and chemokine recognition sites (CRS, underlined), contributing to their different CXCL12 interacting geometries. Receptor residues implicated in CXCL12 β1-strand interactions are underlined. Arrows denote the N-terminal truncation sites of receptors used for molecular dynamics simulations.

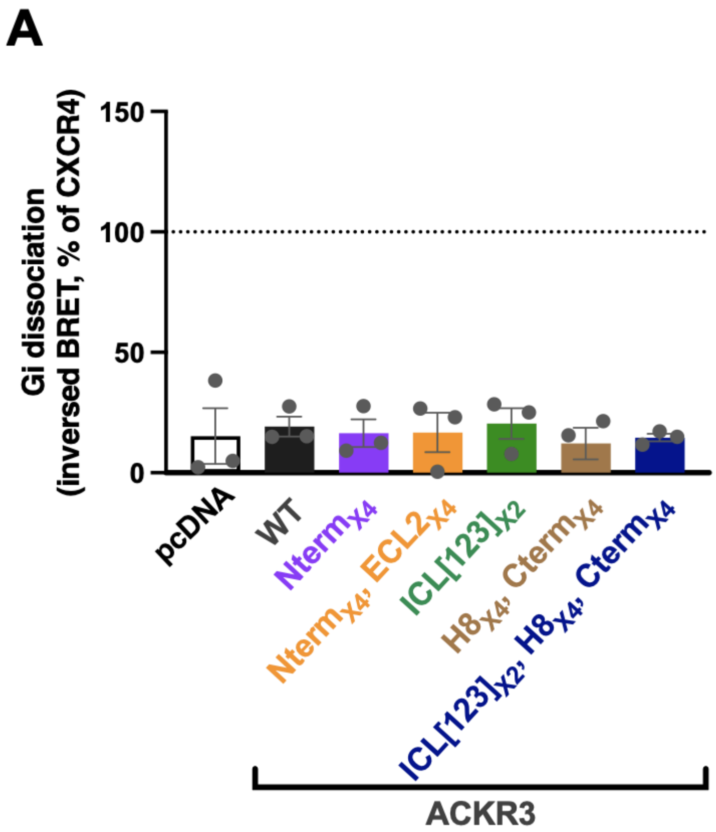


**Figure S3. Chimeric ACKR3 variants containing extracellular and intracellular segments from CXCR2 and CXCR4 fail to activate G proteins.** (**A**) Quantification of Gαi/Gβγ dissociation measured by inverse BRET between Gαi-Nluc and cpVenus-Gβγ. Bar graph showing CXCL12-induced Gαi dissociation for ACKR3 and chimeric mutants containing the canonical sequence substitutions shown in **Fig. 1A**. Responses are expressed as % of the maximal CXCL12-induced Gαi dissociation observed for CXCR4 measured in parallel. Data are derived from the area under the curve (AUC) of net BRET signals over ~10 min following agonist stimulation, corresponding to the representative time courses shown in **Fig. 1D**. (**B**) Surface expression (measured by flow cytometry) of ACKR3 chimeric mutants in the Gαi/Gβγ BRET dissociation experiments. HEK293T cells coexpressing WT ACKR3 or mutants (N-terminally tagged with FLAG) and pIRES Gβ-2A-cpVenus-Gγ2-Gαi-Nluc (same transfectants as used in the BRET experiments) were labeled with anti-FLAG-APC (clone L5) antibody. Cell surface expression was quantified by flow cytometry and expressed as relative fluorescence geometric mean (RFGM). Background fluorescence from pcDNA-transfected cells was subtracted, and values were normalized to WT ACKR3 (100%, dashed line). All data represent mean ± SEM from three independent experiments.

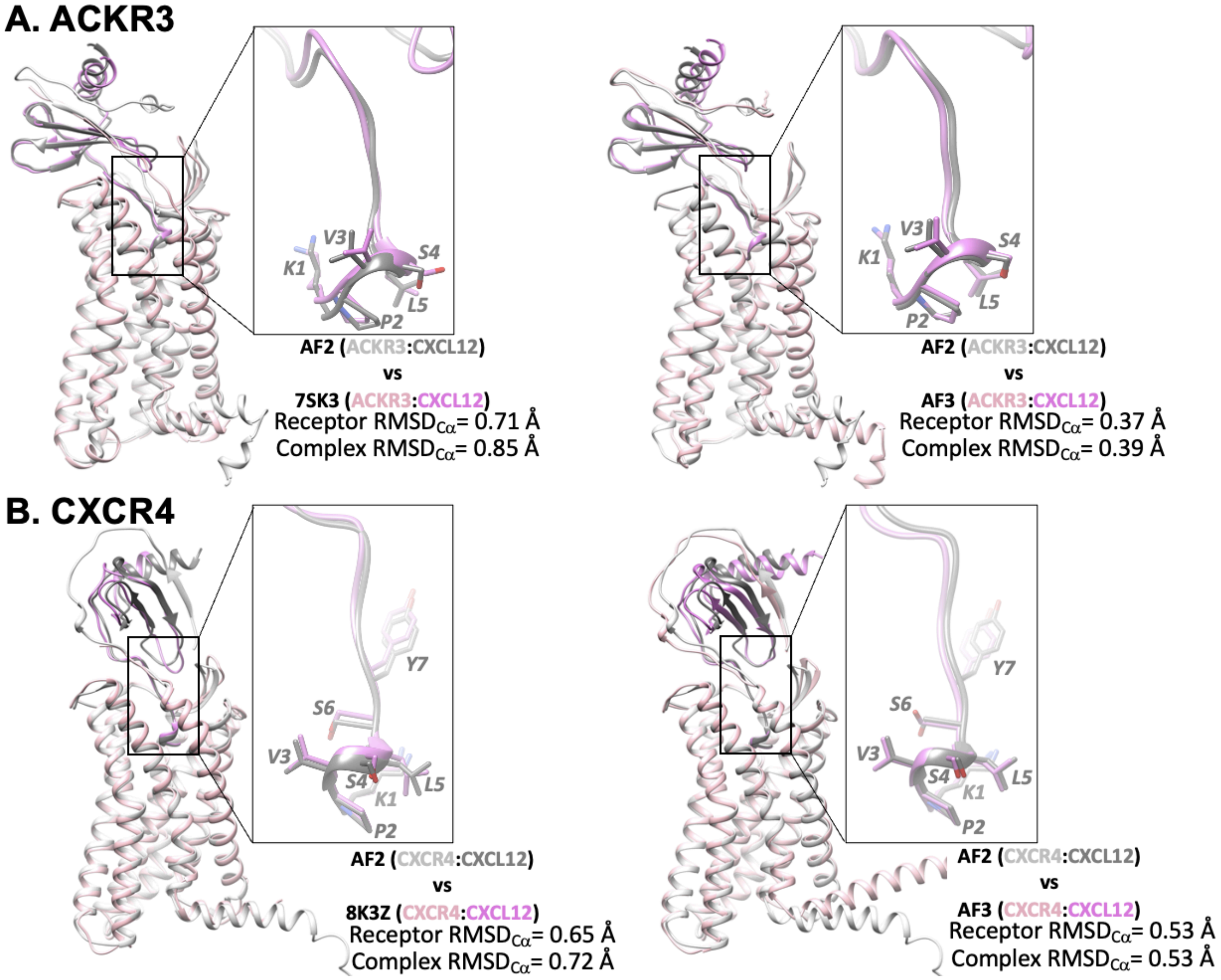


**Figure S4. Agreement of AF2 models with cryo-EM structures and AF3 predictions for ACKR3 and CXCR4 complexes with CXCL12.** Structural alignments of AF2-generated ACKR3:CXCL12 (**A**) and CXCR4:CXCL12 (**B**) complexes used for MD simulations, with their corresponding cryo-EM structures and AF3 models. The alignments reveal nearly identical receptor conformations and CXCL12 binding modes; extended terminal regions were modeled. Models were superimposed with receptors and corresponding $RMSD_{C\alpha}$ values were calculated with PyMOL version 1.20.

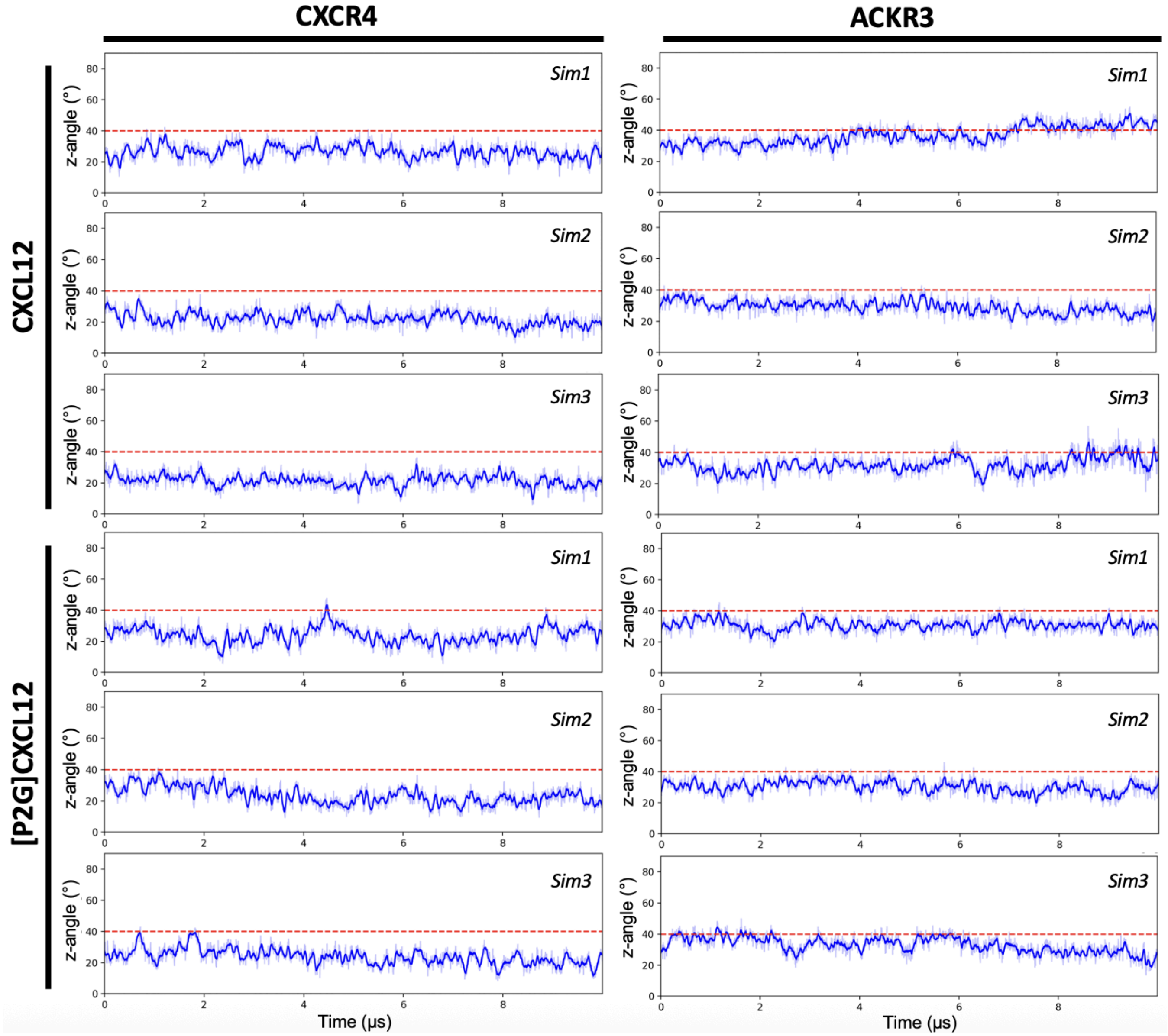


**Figure S5. Time course of receptor tilt angle w.r.t. the z-axis (normal to membrane plane) indicates stable receptor orientations in the lipid bilayer.** The z-angle of the receptor TM backbone is plotted as a function of time. Angles were calculated after centering each frame onto the TMs in the first simulation frame, without alignment. A rolling average (window = 10 frames) is shown to highlight overall trends. The dashed red line marks 40°, shown as a reference threshold.

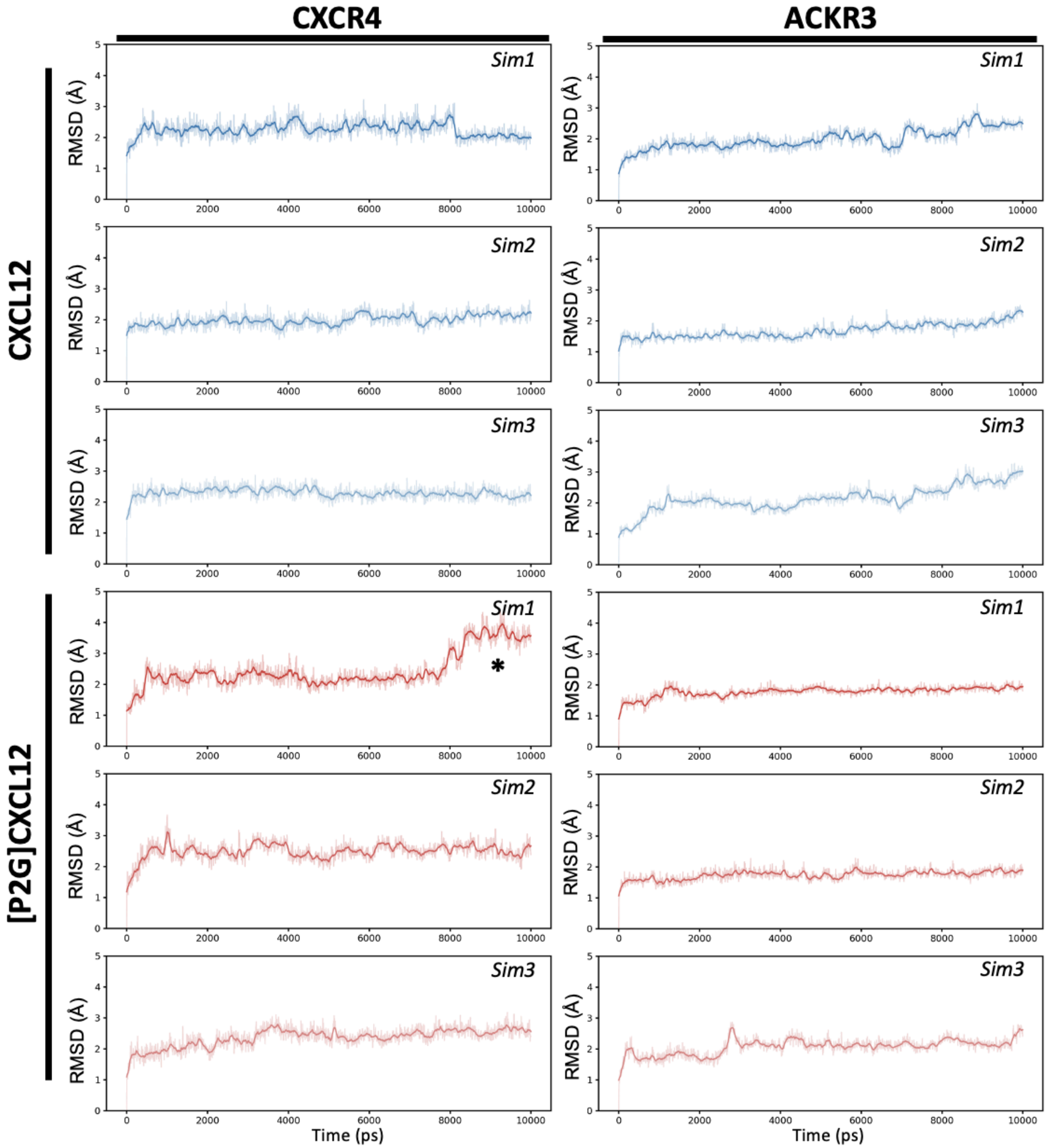


**Figure S6. Time course of receptor TM RMSD indicates overall receptor stability.** RMSD of Cα atoms relative to the starting structure is plotted as a function of time. Each trajectory frame was aligned to the starting structure using receptor TM Cα atoms to remove overall translational and rotational motion, and RMSD was calculated using the same atom selection. A rolling average (window = 20 frames) is shown to emphasize overall conformational trends. The star indicates a time point corresponding to pronounced conformational changes in TM5 and TM6 observed in simulation 1 of CXCR4:[P2G]CXCL12 system (**Fig. S11**).

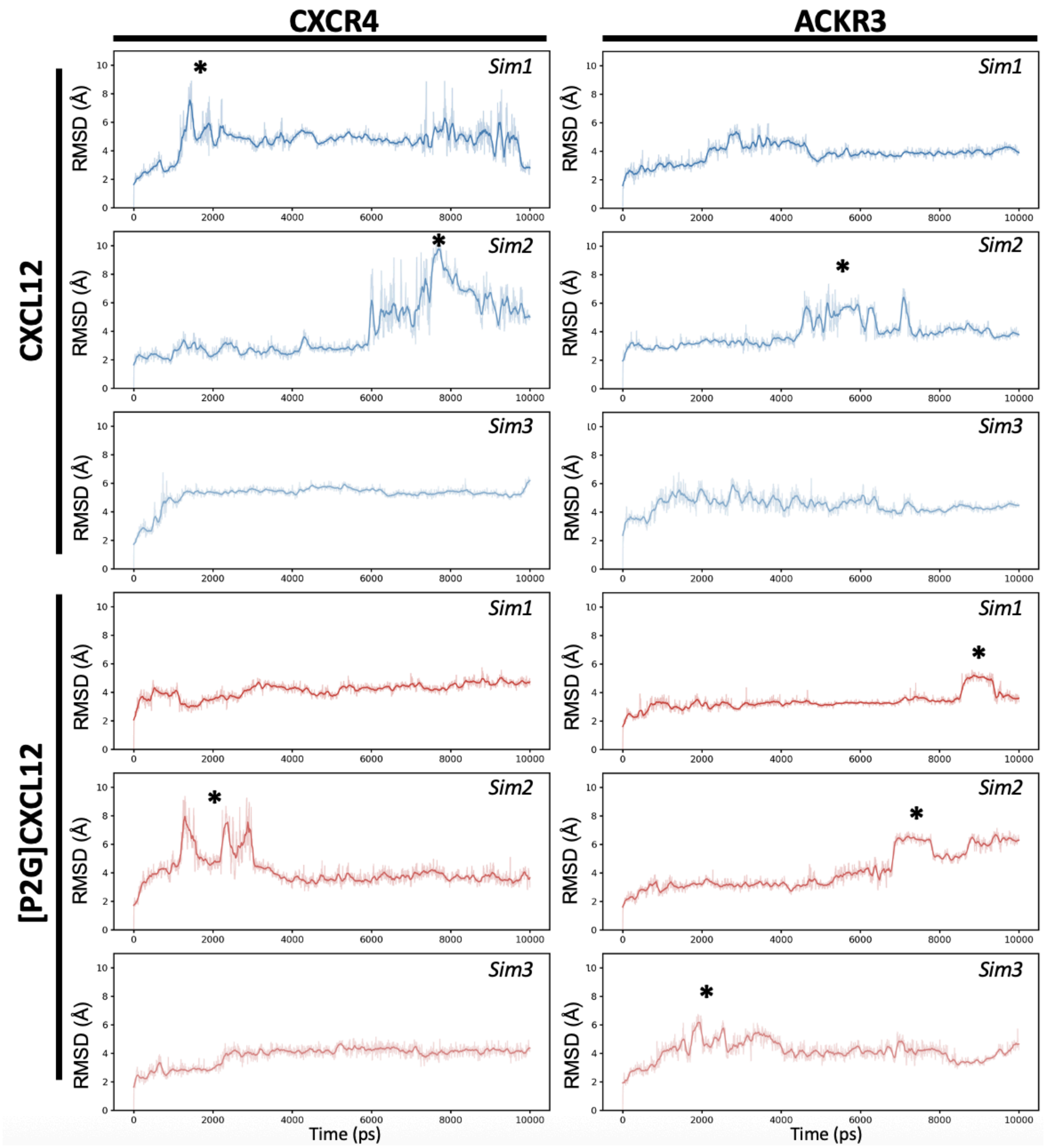


**Figure S7. Time course of the receptor:chemokine RMSD indicates complex stability and transient dissociation of the chemokine β1-strand with receptors.** RMSD of Cα atoms relative to the starting structure is plotted as a function of time. Each trajectory frame was aligned to the starting structure using the receptor:chemokine Cα atoms to remove overall translational and rotational motion, and RMSD was calculated using the same atom selection. A rolling average (window = 20 frames) is shown to emphasize overall conformational trends. Stars indicate stages at which the CXCL12 β1-strand dissociates from receptor N-terminal interaction sites (CRS0.5 in CXCR4 or CRS1 in ACKR3) (**Fig. S12**), resulting in increased flexibility of the receptor N-terminus. This behavior is consistent with the dynamics suggested by cryo-EM structures that show weak or absent density for the receptor N-terminus (1–3). In some trajectories, the β1-strand re-engages the receptor N-terminus, indicating reversible binding dynamics.

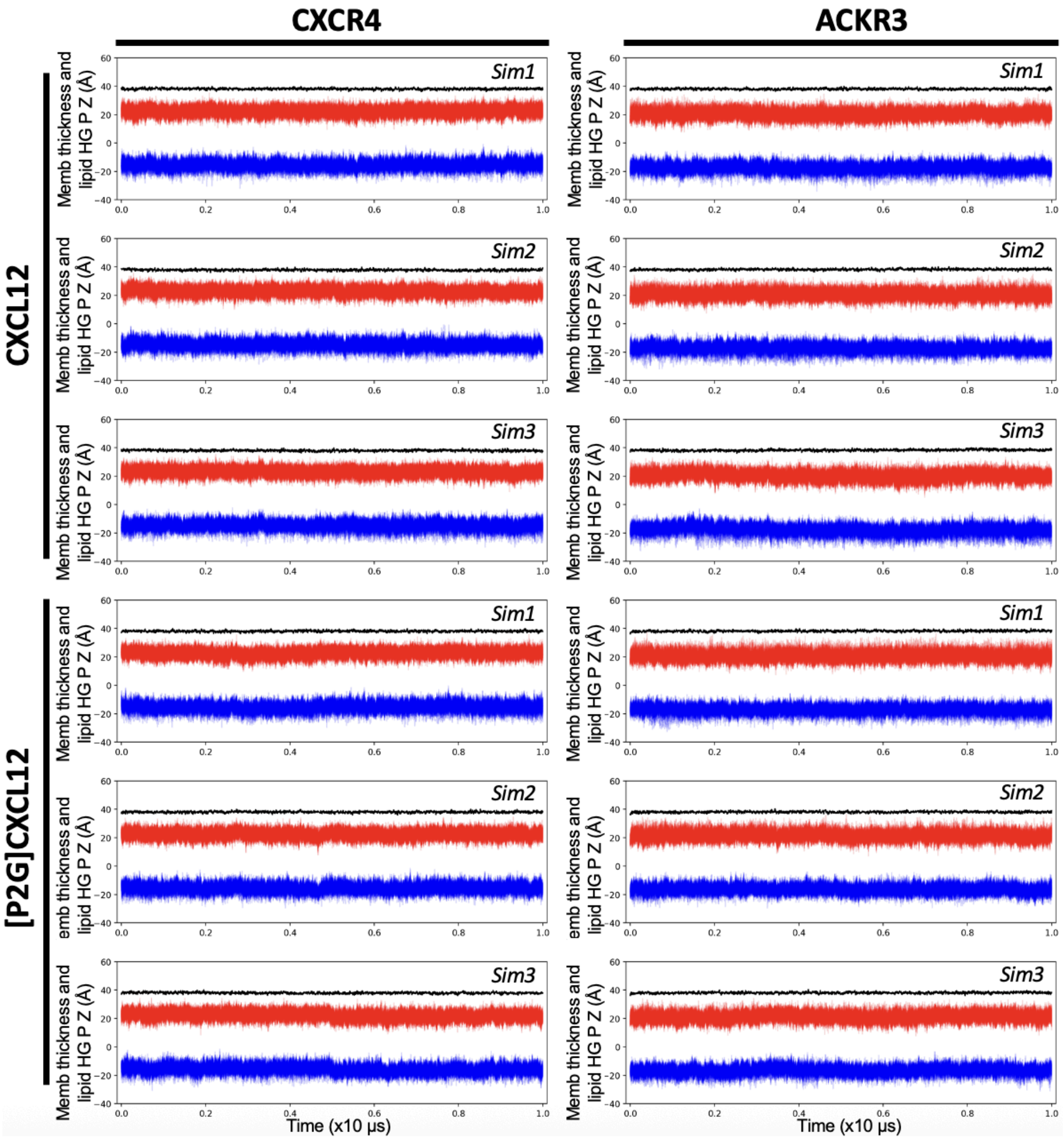


**Figure S8. Time course of membrane thickness and lipid headgroup positions indicates stable membrane organization.** Z-coordinates of lipid phosphate headgroups (P atoms) from individual lipids are shown over time, colored by leaflet assignment (upper leaflet in red, lower leaflet in blue). The membrane thickness, calculated as the distance between the average headgroup positions of the two leaflets, is shown as a black line.

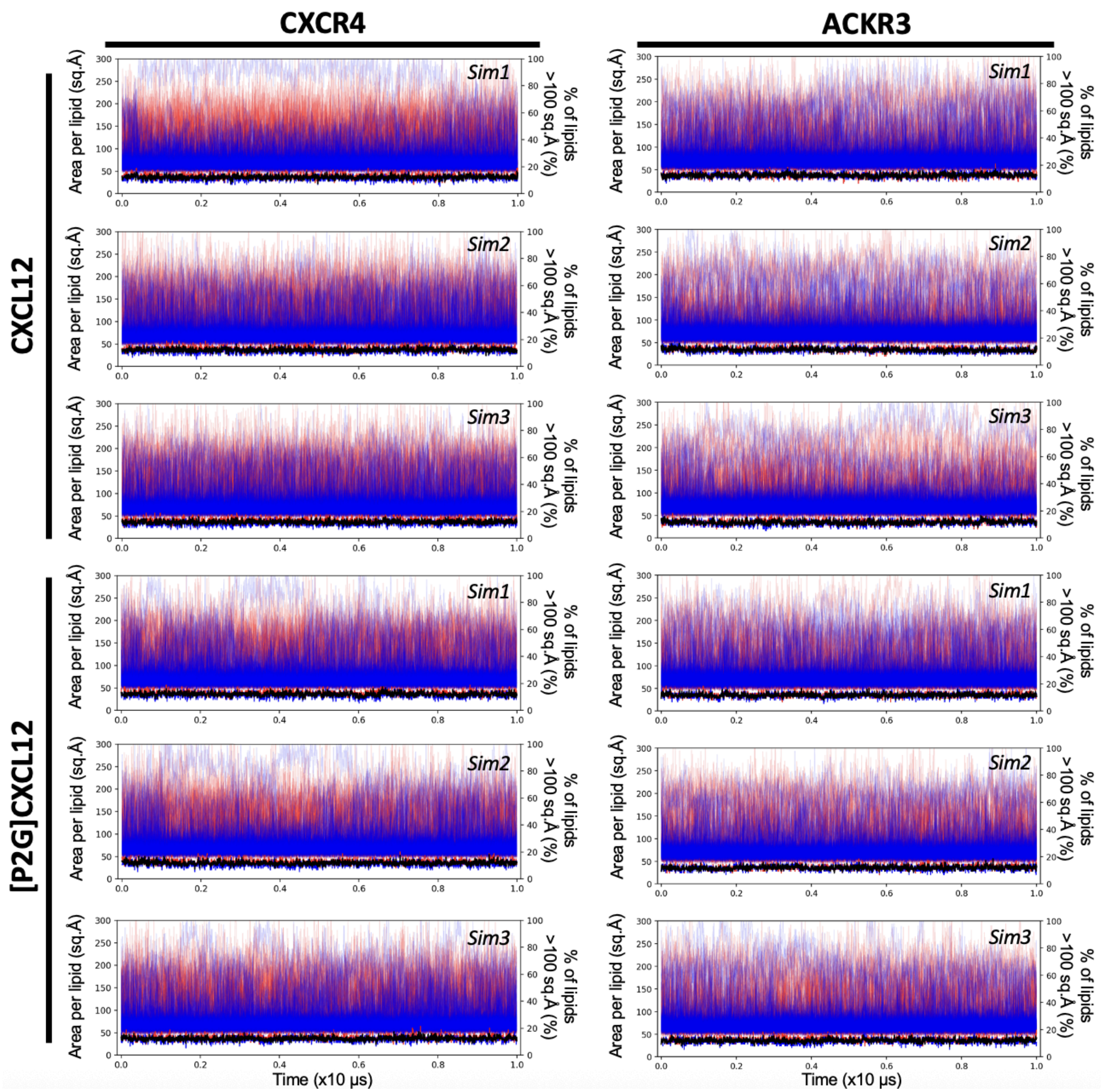


**Figure S9. Time course of area per lipid indicates stable lateral packing of lipids in the membrane.** Area per lipid is shown on the left y-axis (thin traces) for individual lipids, colored by leaflet assignment (upper leaflet, red; lower leaflet, blue). The fraction of lipids with an area greater than 100 Å² is shown on the right y-axis (bold traces) for the upper leaflet (red), lower leaflet (blue), and all lipids combined (black).

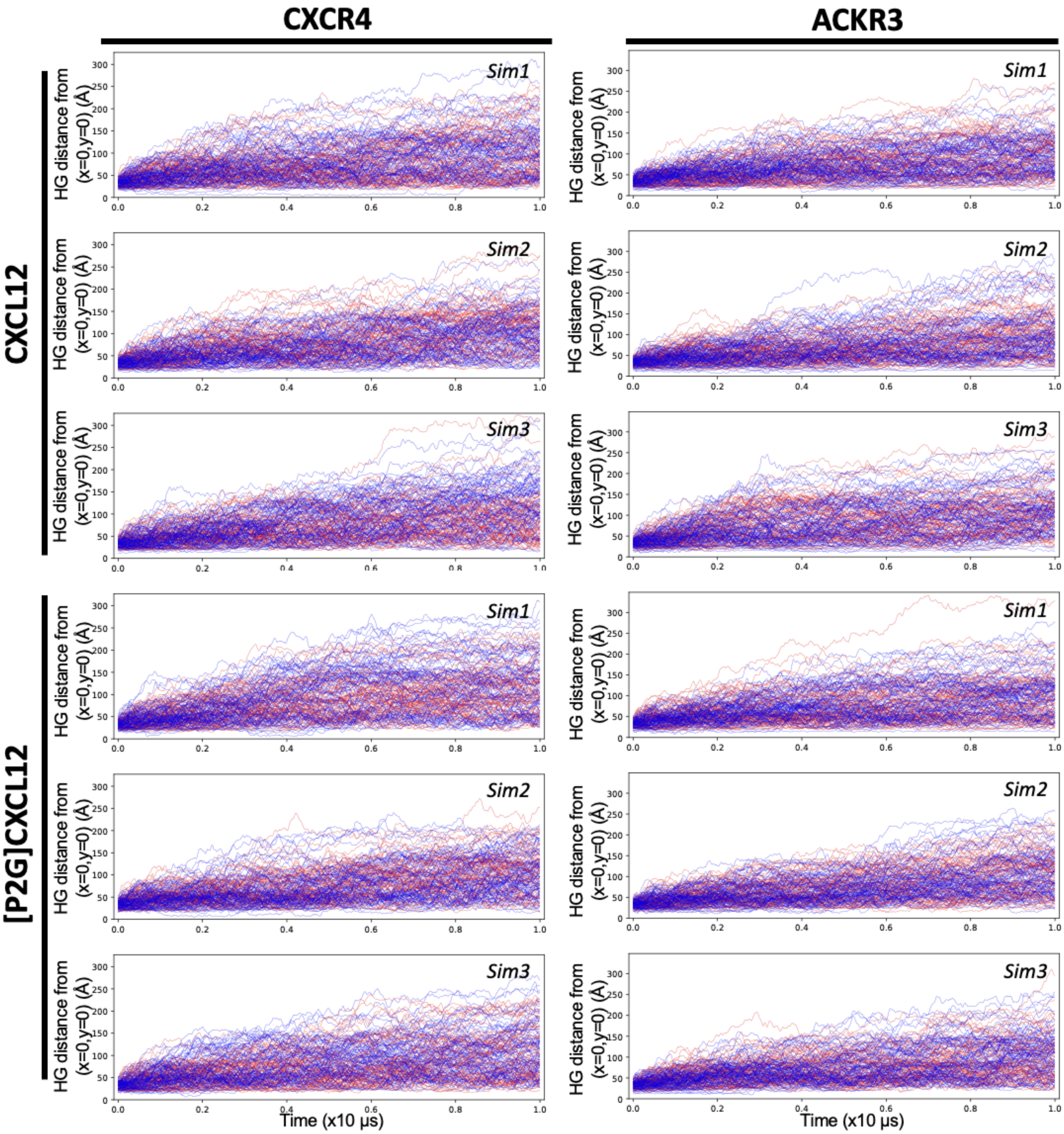


**Figure S10. Lipid headgroup (HG) displacement reflects free lateral diffusion of lipid molecules within a stable membrane.** The radial distance of unwrapped lipid phosphate headgroups from the membrane center (x = 0, y = 0), defined as R = sqrt(x^2 + y^2), is plotted as a function of time. Each trace represents a single lipid headgroup, colored by leaflet assignment (upper leaflet, red; lower leaflet, blue).

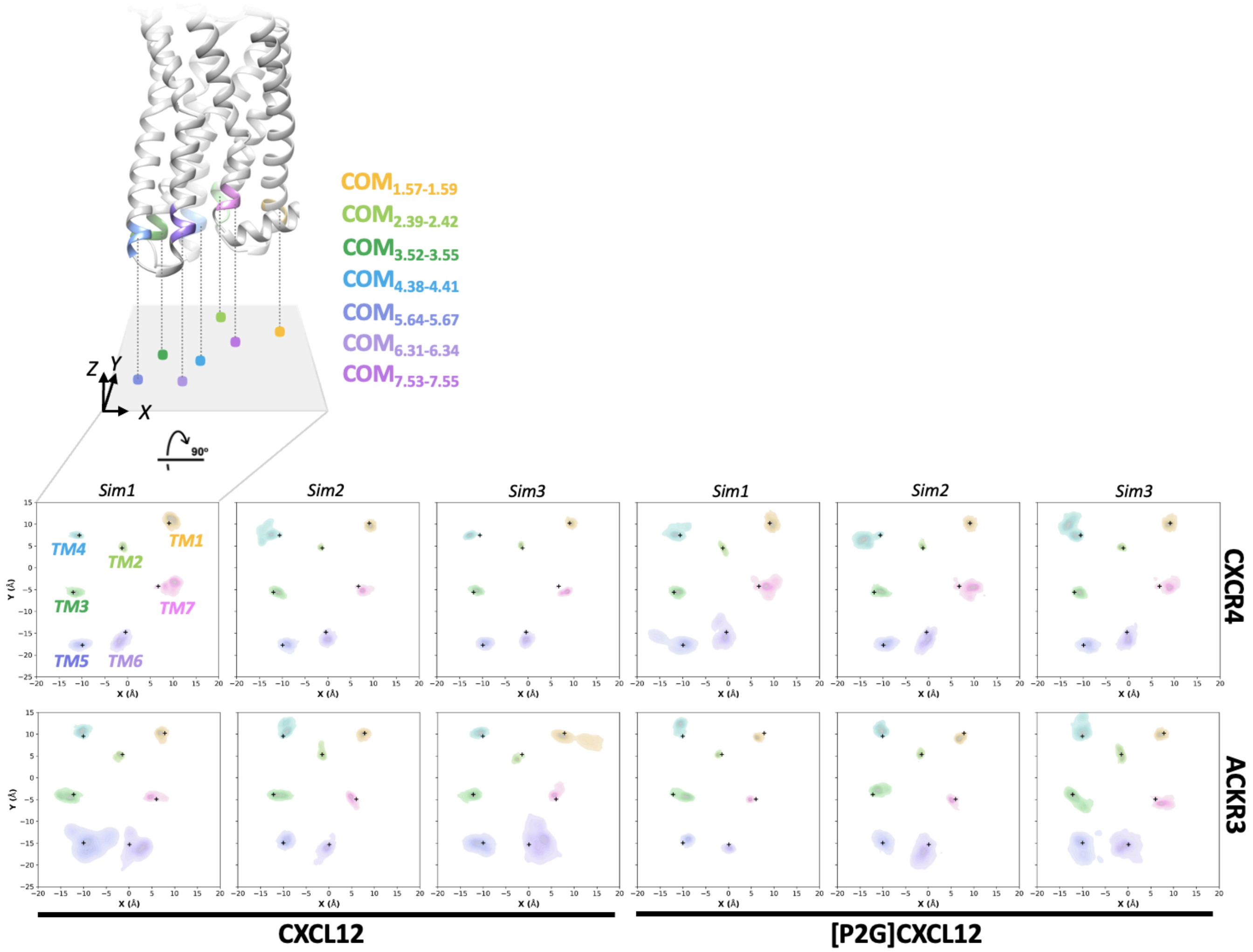


**Figure S11. Replicate MD simulations reveal TM1-7 intracellular conformational dynamics.** Replicate 10 μs trajectories (n=3) aligned to receptor TMs showing conformational dynamics of TM1-7 as center of mass (COM) distributions on the xy-plane. Black '+' indicates original conformations.

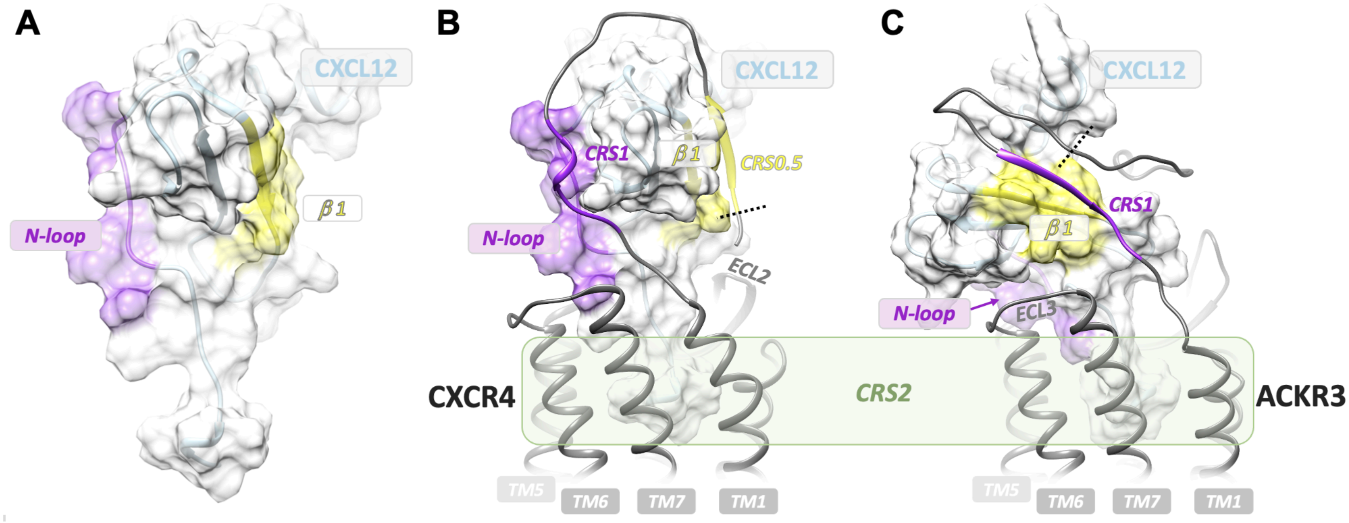


**Figure S12. AF2 models of CXCL12 interactions with CXCR4 and ACKR3 N-termini.** (**A**) CXCL12 structure highlighting the N-loop (residues 12-17) and β1-strand (residues 25-29). (**B**) Model of experimentally verified CXCR4 N-terminal CRS0.5 and CRS1 interactions with CXCL12 β1-strand and N-loop, respectively (4). (**C**) Model of experimentally verified ACKR3 CRS1 interactions with CXCL12 β1-strand (3), showing a distinct binding geometry relative to CXCR4. Dashed lines denote the N-terminal truncation sites of receptors used for molecular dynamics simulations.

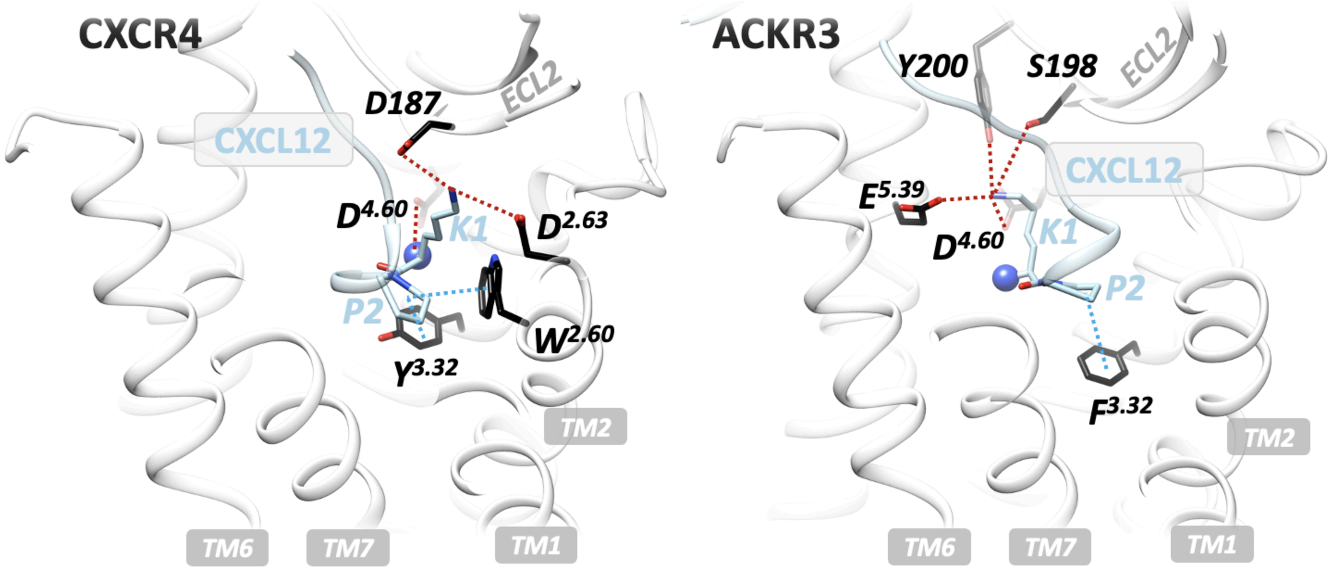


**Figure S13. Distinct interaction networks of CXCL12 Lys1 and Pro2 with CXCR4 and ACKR3.** Cartoon representation of CXCL12 Lys1 and Pro2 interactions within the orthosteric pockets of CXCR4 (PDB 8K3Z) and ACKR3 (PDB 7SK3). In CXCR4, Lys1 forms polar interactions with Asp97$^{2.63}$, Asp171$^{4.60}$, and Asp187$^{ECL2}$, while Pro2 packs against Trp94$^{2.60}$ and Tyr116$^{3.32}$. In ACKR3, the CXCL12 N-terminus adopts a rotated orientation, with Pro2 interacting with Phe124$^{3.32}$ and Lys1 engaging a network involving Asp179$^{4.60}$, Glu213$^{5.39}$, Ser198$^{ECL2}$, and Tyr200$^{ECL2}$. The N-terminal amine of CXCL12 Lys1 is shown as a sphere. Polar and nonpolar interactions are indicated by red and blue dashed lines, respectively.

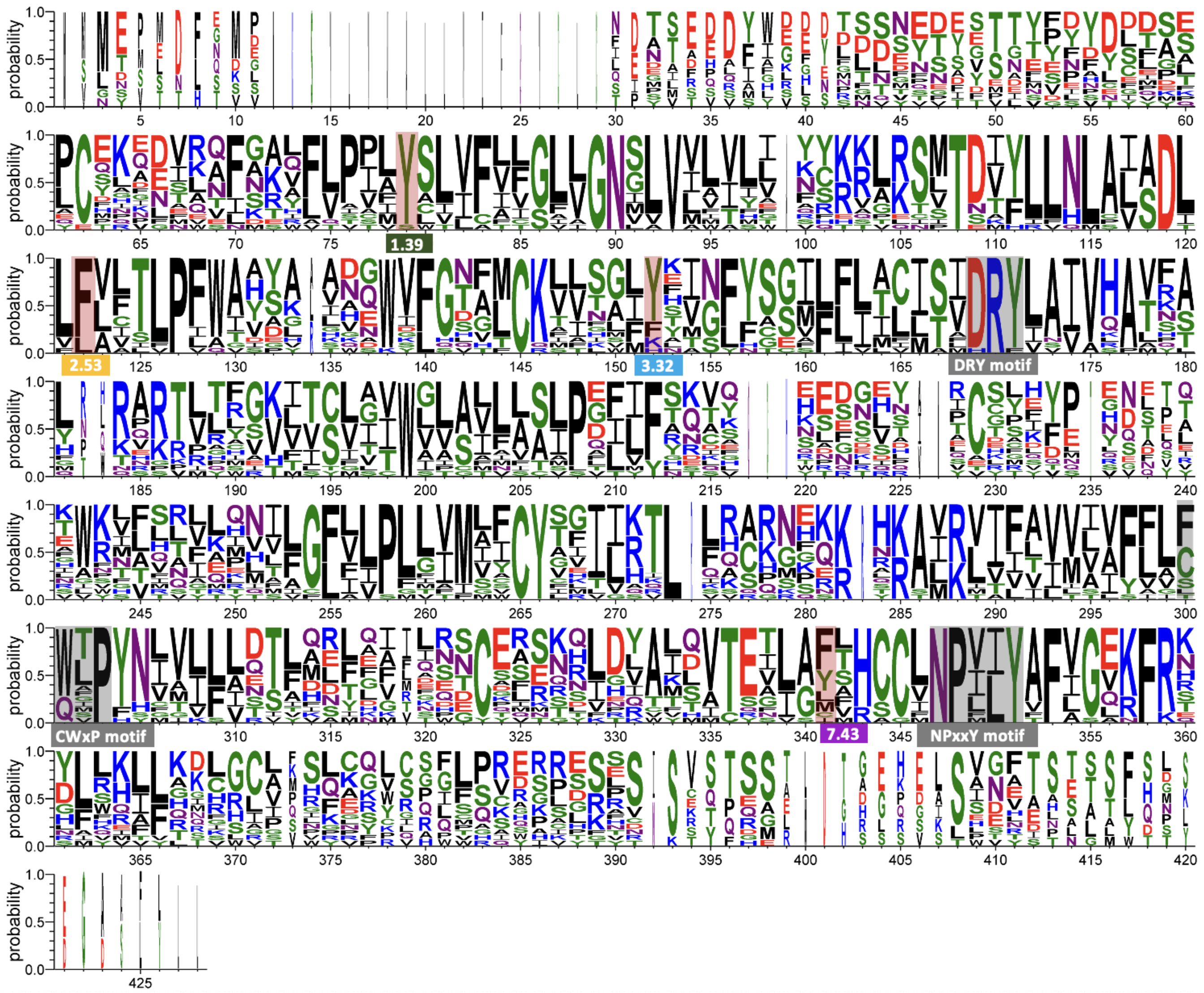

**Figure S14. Conservation of the TM1-TM7-TM2-TM3 aromatic lock in canonical chemokine receptors.** Sequence alignment of CCR1-CCR10, CXCR1-CXCR6, CX3CR1, and XCR1 was generated using Clustal Omega (5) and visualized using WebLogo version 3.9.0 (6). The logo shows strong conservation of residues Tyr$^{1.39}$-Phe$^{7.43}$-Phe$^{2.53}$-Tyr$^{3.32}$ (either Tyr or Phe, labeled as Ballesteros–Weinstein numbering and highlighted in red) forming the aromatic lock. Conserved class A GPCR microswitch motifs, including DRY (TM3), CWxP (TM6), and NPxxY (TM7), are highlighted in gray.

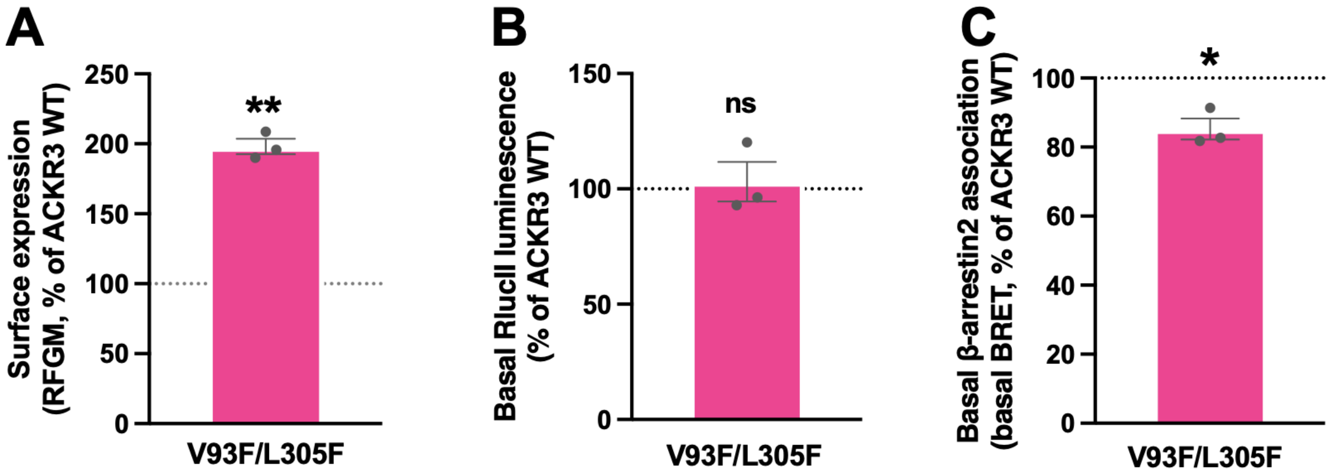


**Figure S15. ACKR3 Val$^{2.53}$Phe/Leu$^{7.43}$Phe (V93F/L305F) shows increased surface expression, relative to WT ACKR3, but decreased basal β-arrestin2 association.** (**A**) Cells were labeled with anti-ACKR3-PE antibody for the detection of the V93F/L305F mutant. Cell surface expression was quantified by flow cytometry and expressed as RFGM. Background fluorescence from pcDNA-transfected cells was subtracted, and values were normalized to WT ACKR3 (100%). (**B**) Total receptor expression was evaluated from the BRET assay by measuring basal RlucII luminescence prior to ligand stimulation and normalized to WT ACKR3 (100%). Similar RlucII signals between WT and mutant indicate comparable receptor expression levels. (**C**) Basal β-arrestin2 association was measured by BRET prior to CXCL12 stimulation and normalized to WT ACKR3 (100%). All data represent mean ± SEM from three independent experiments. Statistical significance was determined using a one-sample t-test comparing mutant to the normalized WT reference; not significant (ns), $p < 0.05$ (*), $p < 0.01$ (**), $p < 0.001$ (***).

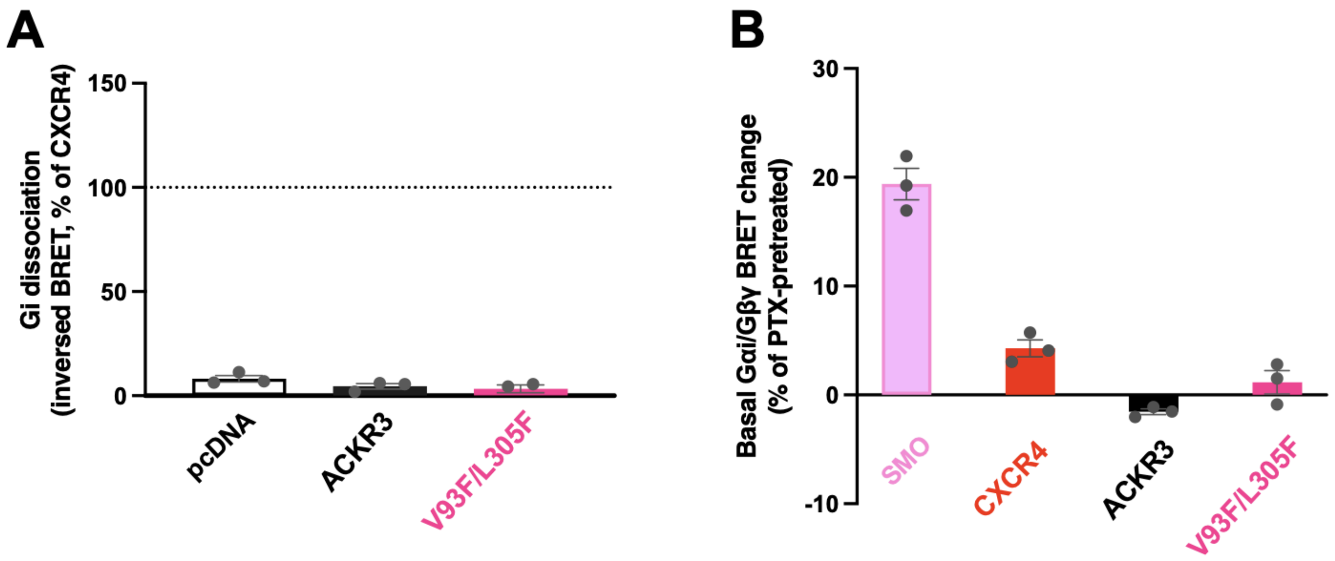


**Figure S16. The ACKR3 V93F/L305F mutant fails to induce Gαi dissociation.** (**A**) Quantification of Gαi/Gβγ dissociation measured by inverse BRET between Gαi-Nluc and cpVenus-Gβγ. Bar graph showing CXCL12-induced Gαi dissociation for ACKR3 and the Val$^{2.53}$Phe/Leu$^{7.43}$Phe (V93F/L305F) mutant. Responses are expressed as % of the maximal CXCL12-induced Gαi dissociation observed for CXCR4 measured in parallel. Data are derived from the AUC of net BRET signals over ~10 min following agonist stimulation. (**B**) Basal Gαi/Gβγ protein dissociation measured by BRET, relative to cells pretreated with pertussis toxin (PTX). Data are presented as mean ± SEM from three independent experiments.

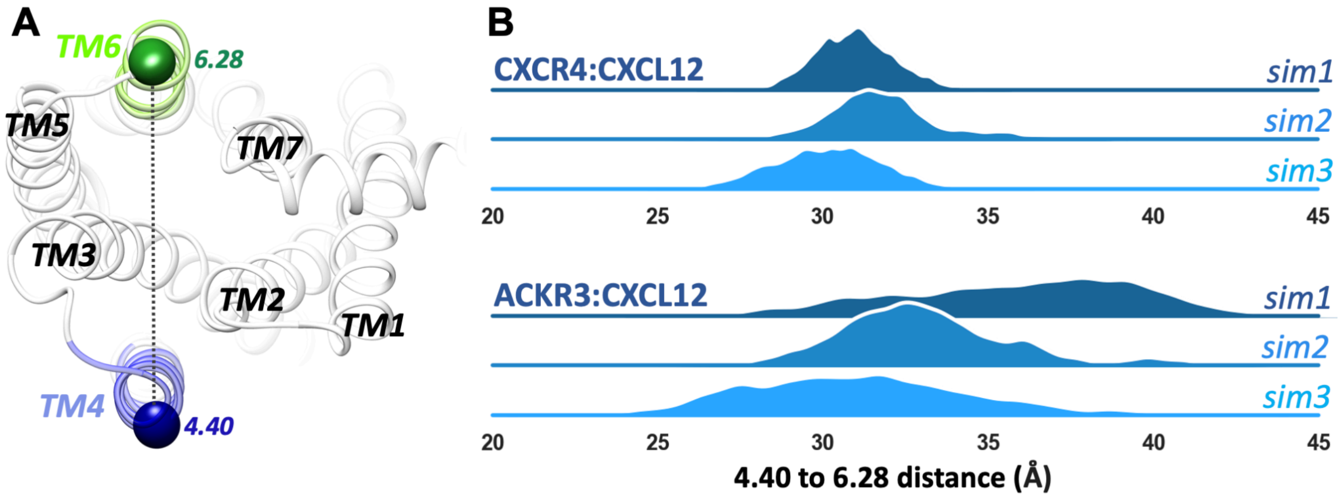


**Figure S17. Measurement of the BW 4.40-6.28 residue distance shows that the ACKR3:CXCL12 complex exhibits greater and more widely distributed TM4-TM6 distances, compared to CXCR4:CXCL12.** (**A**) The intracellular view of ACKR3 TMs. BW residues 4.40 and 6.28, used for ACKR3 labeling in the previous smFRET study (7) were selected for distance measurement. (**B**) Distances between BW residues 4.40 and 6.28 are shown for three independent replicates in CXCR4:CXCL12 vs ACKR3:CXCL12.

**Table S1. Summary of EC50 and Emax values for β-arrestin2 recruitment to receptors and receptor mutants in response to CXCL12 and CXCL12 mutants.**

| Receptors | CXCL12 | EC50 (95% CI) nM | EC50 fold change (vs. WT) | Emax (% of WT CXCL12) | Figures |
|---|---|---|---|---|---|
| CXCR4 | WT | 1.2 (0.3-2.6) | | 100 | 3E |
| CXCR4 | L5F | NA | NA | **6.2 ± 0.9****** | 3E |
| CXCR4 | L5G | NA | NA | **9.5 ± 0.6****** | 3E |
| CXCR4 | L5H | NA | NA | **5.4 ± 0.5****** | 3E |
| CXCR4 | L5N | NA | NA | **12.2 ± 7.5**** | 3E |
| ACKR3 | WT | 18.5 (14.5-23.3) | | 100 | 3G |
| ACKR3 | L5F | 19.3 (14.3-26.4)$^{ns}$ | 1.0 | 98.0 ± 2.1$^{ns}$ | 3G |
| ACKR3 | L5G | 20.2 (17.3-24.3)$^{ns}$ | 1.1 | **90.3 ± 2.1*** | 3G |
| ACKR3 | L5H | 18.5 (14.6-24.0)$^{ns}$ | 1.0 | **90.5 ± 2.3*** | 3G |
| ACKR3 | L5N | **32.9 (25.3-43.6)**** | **1.8** | **85.1 ± 1.5*** | 3G |
| ACKR3 | WT | 8.6 (5.8-12.6) | | 100 | 5F |
| ACKR3 | P2G | 10.4 (7.2-15.1)$^{ns}$ | 1.2 | **89.8 ± 2.2* (*p*=0.0418)** | 5F |
| V93F/L305F | WT | 9.2 (6.6-13.0) | 1.1 | 100 | 5F |
| V93F/L305F | P2G | **19.6 (14.8-26.1)**** | **2.3** | **77.2 ± 2.9* (*p*=0.0158)** | 5F |
| V93F/L305F | WT | 22.2 (14.3-34.9) | | 100 | 5G |
| V93F/L305F | L5F | 25.4 (16.1-40.4)$^{ns}$ | 1.1 | **93.0 ± 1.6*** | 5G |
| V93F/L305F | L5G | **87.9 (72.7-107.1)***** | **4.0** | **55.8 ± 1.1***** | 5G |
| V93F/L305F | L5H | **54.5 (43.0-68.4)**** | **2.5** | **78.8 ± 1.5**** | 5G |
| V93F/L305F | L5N | **92.8 (78.0-110.1)****** | **4.2** | **79.9 ± 1.6**** | 5G |

Dose–response curves were fitted using a four-parameter logistic model in GraphPad Prism version 10.5.0. Emax values were normalized to the maximal response elicited by WT CXCL12 for each receptor construct measured in parallel, and represent mean ± SEM from three independent experiments. NA indicates values that could not be determined due to either absence of detectable agonist activity or failure to obtain a reliable fit (e.g., unstable parameters or excessively wide confidence intervals (CI)). Statistical differences in EC50 values between WT and CXCL12 mutants were assessed using an extra sum-of-squares F test. Emax values were analyzed using a one-sample t-test comparing each mutant to the normalized WT CXCL12 response (100%). Statistical significance is indicated as ns (not significant), *p < 0.05, **p < 0.01, ***p < 0.001, and ****p < 0.0001.

**Table S2. Reagents and supplies.**

| Reagent name | Manufacturer/vendor | Part # |
|---|---|---|
| HEK293T cells | American Type Culture Collection | CRL-3216™ |
| Dulbecco's Modified Eagle Media | Gibco | 11-965-092 |
| Fetal bovine serum | Gibco | A5256801 |
| Trypsin-EDTA (0.05%) | Gibco | 25-300-062 |
| BL21(DE3)pLys competent cells | Invitrogen | C606010 |
| Isopropyl β-D-1-thiogalactopyranoside | Goldbio | I2481C |
| DNaseI | Roche | 10104159001 |
| Ni-NTA agarose resin | Qiagen | 30210 |
| Amicon® centrifugal filter unit | MilliporeSigma | UFC9003 |
| Enterokinase | New England Biolabs | P8070 |
| TransIT®-LT1 transfection reagent | Mirus Bio | MIR 2304 |
| Pertussis toxin | Invitrogen | PHZ1174 |
| Coelenterazine-h | NanoLight Technologies | 301 |
| Prolume Purple | NanoLight Technologies | 369 |
| Accutase | STEMCELL Technologies | 07920 |
| APC-conjugated anti-FLAG antibody | BioLegend | 637307 (clone L5) |
| PE-conjugated anti-ACKR3 antibody | R&D Systems | FAB4227P (clone 11G8) |
